# Individual Dynamics in Stress-Related Pain Responses

**DOI:** 10.64898/2026.01.27.701946

**Authors:** Joren Vyverman, Inge Timmers, Stefanie H Meeuwis, Tom Smeets, Kirsten Hilger

**Affiliations:** Spine, Head and Pain Research Unit Ghent, Department of Rehabilitation Sciences, Faculty of Medicine and Health Sciences, Ghent University, Ghent, Belgium; Pain in Motion International Research Consortium, www.paininmotion.be; Center of Research on Psychological disorders and Somatic diseases (CoRPS), Department of Medical and Clinical Psychology, Tilburg School of Social and Behavioral Sciences, Tilburg University, Tilburg, the Netherlands; Department of Psychology I, Clinical Psychology and Psychotherapy, Würzburg University, Würzburg, Germany; Department of Psychology, Differential Psychology, Personality Psychology and Psychological Diagnostics, Vinzenz Pallotti University Vallendar, Germany

**Keywords:** Individual differences, Maastricht Acute Stress Test, pain sensitivity, stress reactivity, trait pain-related distress

## Abstract

**Background:** Stress and pain are adaptive, bidirectionally related systems, modulated by shared factors. However, how stress reactivity relates to individual differences in pain-related distress and pain sensitivity remains unclear. This preregistered study therefore investigates how physiological reactivity to acute stress relates to trait pain-related distress and stress-induced changes in pain sensitivity, and whether pain-related distress acts as moderator.

**Methods:** Trait pain-related distress was assessed in 148 healthy males using questionnaires. Baseline blood pressure, pulse rate, alpha-amylase, and cortisol were obtained as well as initial heat pain thresholds and tolerances. One group underwent the Maastricht Acute Stress Task, while another group performed the placebo version. Consecutively all stress- and experimental pain indicators were examined again.

**Results:** Trait pain-related distress was not associated with stress reactivity (MAST-induced changes in physiological stress indicators), while stress-induced changes in pain sensitivity showed high individual variability, but were not associated with stress reactivity. Finally, we found preliminary evidence that in individuals with a higher tendency to catastrophize and to fear pain, stronger alpha-amylase increases were associated with larger stress-induced increases in pain threshold (*p-FDR* = .07).

**Conclusion:** Our study illustrates the idiosyncrasy of the complex interplay between trait pain-related distress, physiological stress reactivity, and stress-induced changes in pain sensitivity. While our preliminary results require replication, they suggest that stable individual differences influence the link between stress and pain beyond physiology. This underscores the importance of considering trait differences in future research on stress-pain interactions to, ultimately, better tailoring preventions and treatments for patients with chronic pain.

## 1. Introduction

Stress and pain are bidirectionally related. That is, experiencing acute pain can increase self-perceived stress levels and trigger physiological stress responses (Ulrich-Lai and Herman, 2009). Conversely, exposure to an acute stressor can modulate subsequent pain experience, resulting in either stress-induced increases in pain (i.e., hyperalgesia; Crettaz et al., 2013), decreases in pain (i.e., hypoalgesia; Timmers et al., 2018) or no significant changes in pain (i.e., no response; Gera et al., 2025).

Experiencing pain is an adaptive and protective mechanism. According to the International Association for the Study of Pain (IASP; Raja et al., 2020), pain is defined as an unpleasant sensory and emotional experience underpinned by various neurobiological pathways, including nociceptive transmission, spinal dorsal horn processing and descending modulation (Besson, 1999; Gebhart, 2004). In both research and clinical contexts, pain can be assessed by using quantitative sensory testing (QST; Arendt-Nielsen and Yarnitsky, 2009). Common QST paradigms used in research include pain threshold determination, which is the assessment of the lowest intensity at which a given stimulus is perceived as painful; and pain tolerance determination, which is the assessment of the maximum level of pain a person can tolerate (Orloff, 1979). In addition to these experimental measures, questionnaires are commonly used to evaluate the acute (state) or long-term (trait) emotional, cognitive, and behavioural distress to pain. These affective traits are commonly referred to as ‘pain-related distress’, and research has shown that pain-related distress plays a key role in the development and maintenance of chronic pain and pain-related disability (Crombez et al., 2012).

Similar to pain, experiencing stress is also an adaptive and protective mechanism, enabling the body to respond to potential threats. When a stressor is encountered, the two major stress response systems become activated: the autonomic nervous system (ANS) and the hypothalamic-pituitary-adrenal (HPA) axis. The sympathetic branch of the ANS, also known as the sympatho-adreno-medullary system (SAM), triggers the release of catecholamines in response to stress (Cardinali, 2018). This causes rapid physiological changes, including increased heart rate, blood pressure, and alpha-amylase levels (Rohleder et al., 2004). In contrast, the HPA axis reacts more slowly but produces a longer-lasting effect by releasing cortisol (Kirschbaum et al., 1993).

Although stress and pain follow well-established physiological and neurobiological patterns, there is considerable interindividual variability in how these mechanisms are regulated, experienced, expressed, and in how stress and pain may impact each other. For example, one study observed reduced stress-induced pain inhibition only in individuals with high stress reactivity (i.e., operationalized as changes in self-reported stress), suggesting that acute stress responsiveness may impair pain modulation in certain individuals (Geva and Defrin, 2018). Similarly, recent work by Gera et al. (2025) revealed that stronger stress reactivity (i.e., operationalized as changes in state anxiety levels) was linked to more stress-induced hypoalgesia in people with a lower dispositional tendency for experiencing fear of pain (trait fear of pain) or higher levels of perceived stress in general (Gera et al., 2025). Moreover, Timmers et al. (2018) observed that greater stress-induced decreases in pain sensitivity were associated with higher stress reactivity (i.e., greater cortisol responses) as well as with lower pain-related distress (i.e., trait fear of pain). Together, these findings underscore the complexity of stress-pain interactions. There is evidence of both hypo- and hyperalgesic effects of stress on pain and relatively stable individual differences in pain-related personality traits may shape these interactions. However, it remains unclear whether and how individual differences in pain-related distress are related to stress reactivity, and whether and how such individual differences may impact stress-pain interactions.

Furthermore, research has shown that individual differences in stress responsiveness are common and complex (Kudielka et al., 2009; Rosenblum et al., 2025; Schlotz et al., 2011; Soliemanifar et al., 2018), while various components of the stress response have been implicated in stress-induced changes in pain. Some studies observed evidence that the stress–pain relationship involves the SAM system, but not the HPA axis (e.g., Al’Absi and Petersen, 2003; with effects mediated by blood pressure levels during stress, but not cortisol levels). In contrast, the opposite pattern has also been found (e.g., Timmers et al., 2018; showing associations with changes in cortisol, but not in blood pressure or alpha-amylase). Together, this suggests that the relative contribution of the SAM and HPA axes to pain modulation varies across individuals and contexts. However, a systematic examination of these differences has not yet been conducted.

The aim of this study is to comprehensively examine individual differences in how stress and pain are related in healthy pain-free volunteers, by focusing on relations between trait pain-related distress, physiological stress responses to experimentally induced acute stress (markers of SAM and HPA axes), and stress-induced changes in pain sensitivity. We hypothesized that 1) individuals reporting more trait pain-related distress show greater physiological reactivity to experimentally induced stress, 2) greater stress reactivity is associated with greater stress-induced changes in pain sensitivity, and 3) that lower trait pain-related distress strengthens the association between stress reactivity and stress-induced decreases in pain sensitivity. Comprehensively investigating these complex trait-dependent stress-pain dynamics in a pain-free cohort presents an essential first step for extending our mechanistic understanding to chronic pain populations with the ultimate goal of facilitating the development of personalized therapies and interventions.

## 2. Material and methods

### 2.1 Preregistration

The full research project is accessible on the Open Science Framework (see: https://osf.io/zh3ak/). The main objectives of this study were preregistered (see: #56126 | AsPredicted). Note that to provide more comprehensive insights into stress-pain interactions (rather than a secondary focus on other personality traits), the present study differs from this preregistration in the additional inclusion of secondary analyses and the consideration of pain thresholds and tolerances as outcome measures beyond those already preregistered.

### 2.2 Study design and objectives

This study is part of a larger cross-sectional research project (i.e., Fear of Pain, Stress and Individual Differences in affect and cognition; FOPSID, Hilger et al., 2023) and is reported in accordance with the STROBE checklist for cross-sectional studies (Von Elm et al., 2007). The objectives of this study are: (**O1**) to examine the association between individual differences in stress reactivity and trait pain-related distress; (**O2**) to assess the association between individual differences in stress reactivity and changes in pain sensitivity following acute stress; and (**O3**) to determine whether trait pain-related distress moderates the relationship between stress reactivity and stress-induced changes in pain sensitivity.

### 2.3 Participants

One hundred and fifty-two participants volunteered in this study and were assigned randomly to the experimental stress group or the no-stress control group. Four participants were excluded from further analyses due to dropout, resulting in a final sample of 148 participants. G*power analyses (version 3.1; exact test family, bivariate normal model for correlation, based on O1 and its corresponding preregistered hypothesis) considering small effect sizes (Pearson correlation coefficient) of .25, α of .05 and a power of .95 resulted in a required sample size of 168 participants (Faul et al., 2009). However, due to practical restrictions during the COVID-19 pandemic, it was not feasible to achieve the originally planned sample size, resulting in a slightly diminished power of .93. Participants were recruited using different channels (e.g., social media, flyers, University platform). Only males between 18-35 years old, with a BMI between 18-30 kg/m^2^ and with sufficient German language abilities were included. Interested volunteers were excluded when one of the follow criteria were met: studying psychology beyond the second semester, being left-handed, regular intake of medication affecting the central nervous system and psychological functions (e.g., methylphenidate, antidepressants), drug consumption, frequent smoking behaviour (>15 cigarettes per day), acute suffering from a current or past chronic pain condition (e.g., headache), seeking medical treatment in the past six months for an episode of severe pain, current or history of psychological, neurological, cardiovascular, respiratory or endocrinological disease, and suffering from neurodermatitis. A telephone interview was conducted to confirm that all inclusion criteria were met. Participants were rewarded with 50€ for their participation. All procedures were conducted according to the declaration of Helsinki and approved by the ethics committee of the Psychological Institute of the University of Würzburg (GZEK 2020-18).

### 2.4 Study procedure

An overview of the study procedure is illustrated in **Figure 1**. While the larger FOPSID project consisted of three sessions conducted on three separate days (comprising also a Virtual Reality task on day two and an electroencephalogram on day three), in this study, only data from the first two sessions were included before the Virtual Reality part started.

**Figure 1.**
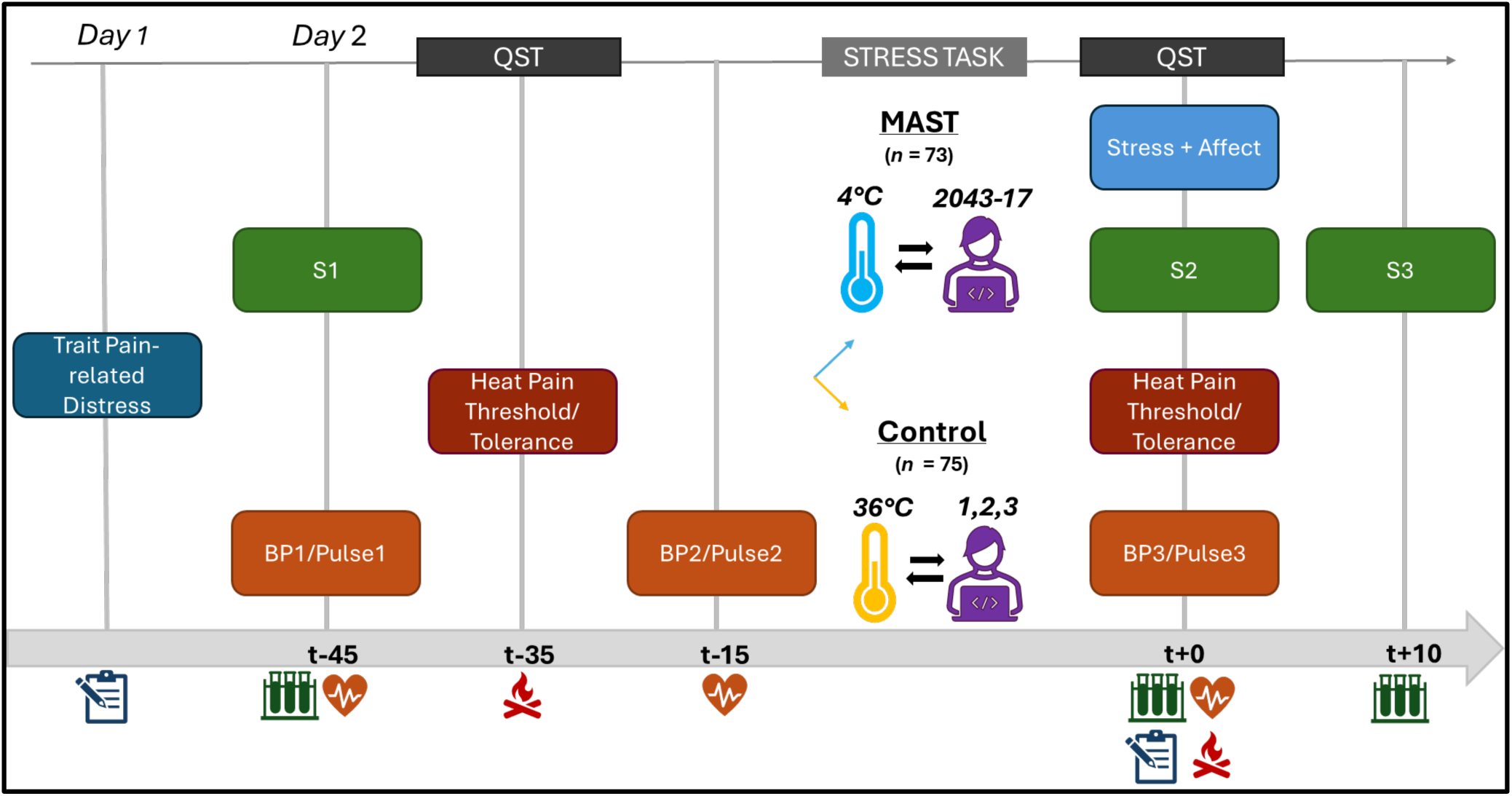
Overview of the experimental paradigm. BP: Blood Pressure, MAST: Maastricht Acute Stress Task, S: Salivary (alpha-amylase and cortisol) Sample, QST: Quantitative Sensory Testing. Participants completed trait pain-related distress questionnaires, followed by baseline assessments of blood pressure, pulse, alpha-amylase, cortisol and heat pain threshold/tolerance. Immediately before the stressor, blood pressure and pulse were reassessed. Participants were then randomly assigned to either the MAST or the control condition. After the stress task, subjective ratings (stress, pain, unpleasantness, affect due to MAST), blood pressure, pulse, alpha-amylase/cortisol, and heat pain threshold/tolerance were obtained. A final alpha-amylase/cortisol sample was collected 10 minutes post-task.

On the *first day*, participants were informed about the study details, they were assured that they could withdraw at any time without consequences, and the informed consent form was signed. Then, trait questionnaires, including the German version of the Fear of Pain Questionnaire (FPQ-III; Baum et al., 2013; McNeil and Rainwater, 1998), the German version of the Pain Catastrophizing Scale (PCS; Meyer et al., 2008; Sullivan et al., 1995) and the general population adaptation of the Tampa Scale of Kinesiophobia (TSK-G; Houben et al., 2005) were filled out.

Before starting the *second day*, participants were instructed to be awake at least four hours before the start of the experiment, to abstain from consuming alcohol at least 12 hours and caffeinated drinks two hours before testing, and to avoid the usage of pain medication on the day of testing. Testing days ran between 11:00 a.m. and 8:00 p.m. to minimize morning fluctuations in cortisol levels. To start, a demographic questionnaire was filled out. Afterwards, baseline measurements were taken, including blood pressure (BP1), pulse (Pulse1) and a salivary alpha-amylase and cortisol sample (S1). Heat pain thresholds (HPTh) and heat pain tolerances (HPTo) were assessed prior to the initiation of the Maastricht Acute Stress Task (MAST; Smeets et al., 2012) or its control condition. Blood pressure and pulse were measured again before the stressor (BP2, Pulse2) and immediately following its completion (BP3, Pulse3). Salivary samples were collected immediately after the stressor (S2) and 10 minutes after the end of the stressor (S3). Subjective stress levels, pain levels, and unpleasantness due to the MAST were evaluated using a Visual Analog Scale (VAS), and positive and negative affect were assessed with the German version of the Positive and Negative Affect Schedule (PANAS; Breyer and Bluemke, 2016; Watson et al., 1988), both administered immediately following the stressor. HPTh and HPTo were reevaluated after completion of the stressor.

### 2.5 Stress induction

The MAST is a reliable, validated acute stress induction task (Smeets et al., 2012). It consists of a five-minute preparation/instruction phase and a 10-minute acute stress phase including physical, cognitive and social stressors during which physical and cognitive stressors altered. The physical stressor consisted of immersing the hand into a cold-water tub (JULABO Labortechnik GmbH, Seelbach, Germany) at 4°C for 60 or 90 seconds per trial, while the cognitive stressor consisted of counting backwards in steps of 17 starting from 2043 as quickly and accurately as possible for 45, 60 or 90 seconds per trial. The duration and order of the trials were fixed; however, participants were told the duration of each trial would be randomly chosen by the computer to increase unpredictability of the stress duration (maximally 90 seconds). As social stressor, participants received negative feedback during counting, and participants thought they were videotaped for analyses of facial expressions. The control-MAST consisted of immersing the hand in the water bath at 36°C, an elementary counting task (counting from one to 25), and no social stressor (Smeets et al., 2012).

### 2.6 Stress assessments

Blood pressure and pulse were measured with a blood pressure monitor (Sanitas SBM 21 upper arm blood pressure monitor; Hans Dinslage GmbH, Germany). Mean arterial pressure (MAP) was calculated in order to obtain one average blood pressure value ([MAP = DBP + 0.412 (SBP – DBP)]; Meaney et al., 2000). Reactivity of the SAM axis was calculated as the change in MAP and pulse from immediately pre-stressor to immediately post-stressor (Δ MAP/Pulse: MAP3 – MAP2/Pulse3 – Pulse2). Salivary alpha-amylase and cortisol were collected using a synthetic salivate (Sarstedt, Germany). Participants were instructed to place the cotton in their mouths without manual contact for 60 seconds. Afterwards, the sample was immediately stored at -20°C. Concentrations of alpha-amylase were measured by an enzyme kinetic method (Rohleder et al., 2006). Cortisol levels were determined using a commercially available chemiluminescence immunoassay kit with high sensitivity (IBL International, Hamburg, Germany). Mean intra- and inter-assay coefficients of variation were less than 7%. Changes in alpha-amylase were calculated as the change from baseline to immediately after the stressor (Δ alpha-amylase: S2 – S1), whereas HPA axis reactivity was quantified as the change in cortisol levels from baseline to 10 minutes post-stressor (Δ cortisol: S3 – S1). Furthermore, the area under the curve with respect to the increase (AUCi) was calculated as a total value for cortisol increases in response to the MAST (Pruessner et al., 2003).

Finally, a 0-100 VAS was used to also measure the subjective experience of the MAST by rating how stressful, painful and unpleasant the MAST procedure was (0 = “not at all”; 100 = “extremely”). Positive and negative affect were measured using the PANAS (Cronbach’s α = .86) (Breyer and Bluemke, 2016; Watson et al., 1988), which consists of 10 questions concerning positive affect (e.g., enthusiastic, excited), and 10 questions concerning negative affect (e.g., distressed, afraid). Each question was scored between one and five on a Likert scale (1 = “not at all”; 5 = “extremely”).

### 2.7 Trait pain-related distress

#### 2.7.1 Pain Catastrophizing Scale

To obtain individuals’ tendency to have catastrophic thoughts regarding pain, the PCS questionnaire (Cronbach’s α = .92) was used. It consists of 13 items, scored on a five-point Likert scale ranging from 0 = “not at all” to 4 = “all the time”. Total scores range from 0 to 52, with higher scores indicating a higher tendency to have catastrophic thoughts regarding pain (Meyer et al., 2008; Sullivan et al., 1995).

#### 2.7.2 Tampa Scale of Kinesiophobia

Fear of movement or injury was assessed with the TSK-G questionnaire (Cronbach’s α = .72), containing 17 items, scored on a four-point Likert scale, ranging from 1 = “strongly disagree” to 4 = “strongly agree”. Total scores range from 17 to 68, with higher scores indicating severe fear of movement or injury (Houben et al., 2005).

#### 2.7.3 Fear of Pain Questionnaire

To measure trait fear of pain, the FPQ-III (Cronbach’s α = .90) uses 30 items, scored on a five-point Likert scale ranging from 1 = “not at all” to 5 = “extreme”. Total scores range from 30 to 150, with higher scores indicating stronger fear of pain (Baum et al., 2013; McNeil and Rainwater, 1998).

### 2.8 Experimental outcomes of pain

HPTh and HPTo were assessed using a Peltier-type thermode (Somedic MSA thermal stimulator; Somedic Sales B, Hörby, Sweden), which was attached to the participant’s left volar forearm. For HPTh, participants pressed a response button when the temperature became painful; for HPTo, they pressed it at the maximum temperature they were willing to tolerate. Each measure was derived from five trials, with temperatures increasing from 32 °C at 1 °C/s, interstimulus intervals of 4–6 s, and a hardware-limited maximum of 49 °C. Analyses used the mean of the final three trials for both HPTh and HPTo. Stress-induced changes were calculated as the pre-stressor minus post-stressor values (ΔHPTh/HPTo = HPTh/HPTopre-stressor – HPTh/HPTopost-stressor).

### 2.9 Data and statistical analysis

All statistical analyses were conducted in R (Version 4.5.1, R Core Team, 2025). Normality and distribution of outcome variables were assessed using the Shapiro-Wilk test, as well as through visual inspection. Outliers were identified as values exceeding ±3 standard deviations (SD) from the mean (M) and were winsorized to the respective M ±3 SD threshold. Descriptive statistics were compared between the stress group and the control group using independent samples *t*-tests or Wilcoxon rank-sum tests, depending on the normality of the data. Cortisol analyses included only participants who adhered to the instructions, whose baseline cortisol levels were below 15 nmol/L (as higher values were interpreted as indicative of not adhering to the instructions or reflecting anticipatory stress), and whose baseline cortisol levels did not exceed ±3 SD (Smeets et al., 2019). Additionally, cortisol responders (increase >1.5 nmol/l; Miller et al., 2013) in the control group were excluded from all further analyses of cortisol reactivity to account for unintended physiological reactivity. Due to the typical right-skewed distribution of cortisol and alpha-amylase values, both variables were log-transformed prior to analysis.

To evaluate the effectiveness of the stress manipulation, group differences over time in physiological stress markers were examined with linear mixed-effects models (LMM) and restricted maximum likelihood (REML) in R (lme4, lmerTest, emmeans, effectsize packages in R). Models included fixed effects of group (stress vs. control), time (baseline, pre-stressor, post-stressor), and interaction with random intercepts. Model assumptions and overall model fit were evaluated using visual inspection and statistical diagnostic procedures. Fixed effects were assessed with Type III *F*-tests (Satterthwaite). Effect sizes (*η²ₚ*) are reported and post-hoc pairwise comparisons of estimated marginal means were conducted to further unravel the observed effects. The Holm–Bonferroni method was used to control the family-wise error rate within each set of comparisons, and adjusted *p*-values are reported alongside uncorrected *p*-values. Post-MAST subjective stress, pain, unpleasantness, and positive and negative affect after the MAST were compared between groups using independent *t*-tests or Wilcoxon rank-sum tests, depending on normality.

Simple bivariate associations between physiological stress reactivity (Δ MAP, Δ pulse, Δ salivary alpha-amylase, Δ cortisol) and trait pain-related distress (PCS, TSK, PFQ sum scores) (**O1**), as well as (stress-induced changes in) pain sensitivity (Δ HPTh, Δ HPTo) (**O2**), were examined using Pearson or Spearman correlations, depending on data normality. Changes in experimental pain outcomes and time-by-group interactions were assessed with LMMs.

Lastly, to investigate whether trait pain-related distress influences the strength of the relationship between stress reactivity and stress-induced changes in pain sensitivity, moderation analyses were performed in the stress group (**O3**). For each outcome variable (Δ HPTh, Δ HPTo), four markers of stress reactivity (Δ MAP, Δ pulse, Δ salivary alpha-amylase, Δ cortisol) in combination with trait pain-related distress variables as moderators (PCS, TSK, PFQ sum scores) were tested. To facilitate interpretation of interaction effects (Aiken et al., 1991), all predictors and moderators were mean-centred before model estimation. Each model included main effects of predictor and moderator, and the interaction term. In all analyses, model assumptions were checked systematically, using the Shapiro-Wilk test for normality, the Breusch-Pagan test for homoscedasticity, variance inflation factors for multicollinearity, and Cook’s distance for influential observations. When statistical assumptions were met, inference was based on the standard Ordinary Least Squares (OLS) estimation, otherwise, non-parametric bootstrapping (5,000 resamples, percentile 95% confidence interval; CI) was used to re-estimate the interaction effect. Model coefficients (*B* and *β*), standard errors (SE), *t*-statistics, 95% CI, *p*-values and explained variance effect sizes (*R²*, *ΔR²*, *f²*) were reported. Multiple comparisons were controlled using the False Discovery Rate (FDR; Benjamini–Hochberg), which was applied within each set of related tests, examining one hypothesis. Significance was defined by *p* < .05.

Exploratively, all objectives were tested in cortisol responders, and comparisons of trait pain-related distress and stress reactivity were made between three subgroups classified based on the stress-induced pain response: Stress-induced hyperalgesia (i.e., decrease in pain threshold >1°C; SI-hyper) group, stress-induced hypoalgesia (i.e., increase in pain threshold >1°C; SI-hypo) group and stress-induced non-response (i.e., no change >1°C; SI-NR) group (Gera et al., 2025) using one-way ANOVA or Kruskal-Wallis when normality assumptions were violated.

## 3 Results

### 3.1 Participant characteristics

Participant characteristics are represented in **Table 1**, including 73 participants in the experimental stress group and 75 participants in the control group. Eight participants exceeded the ±3 SD threshold and/or demonstrated cortisol values at baseline >15 nmol/l. Those participants (*n* = 5 in the stress group, *n* = 3 in the control group) were excluded from the primary analyses regarding cortisol. Furthermore, eight participants in the control group showed an increase of ≥1.5 nmol/l due to the placebo-stress induction and were therefore also excluded from the primary analyses involving cortisol and both groups of participants (cortisol analyses: stress group, *n* = 68; control group, *n* = 64).

**Table 1.** Participant characteristics. . Values are expressed as mean (standard deviation). AUCi: Area Under the Curve with respect to the increase, HPTh: Heat Pain Threshold, HPTo: Heat Pain Tolerance, MAP: Mean Arterial Pressure, MAST: Maastricht Acute Stress Task.

|  | **Total**<br>**(n = 148)** | **Stress group**<br>**(n = 73)** | **Control**<br>**group**<br>**(n = 75)** | **Group**<br>**comparison** |
| --- | --- | --- | --- | --- |
| Age (years) | 24.73 (3.99) | 24.47 (3.76) | 24.99 (4.20) | $p = .501$ |
| Pain catastrophizing (PCS; 0-52) | 17.84 (7.60) | 18 (7.28) | 17.68 (7.94) | $p = .697$ |
| Kinesiophobia (TSK-G; 17-68) | 34.82 (5.60) | 34.89 (5.06) | 34.75 (6.11) | $p = .880$ |
| Fear of pain (FPQ-III; 30-150) | 70.49 (14.96) | 72.19 (13.49) | 68.83 (16.17) | $p = .171$ |
| Baseline MAP (mmHg) | 102.38 (8.45) | 101.70 (8.92) | 103.04 (7.97) | $p = .611$ |
| Baseline pulse (beats/minute) | 81.57 (21.87) | 82.08 (21.52) | 81.08 (22.35) | $p = .444$ |
| Baseline alpha-amylase (U/mL) | 101.24 (108.53) | 103.87 (113.03) | 98.64 (104.61) | $p = .982$ |
| Baseline cortisol (nmol/l) | 5.65 (4.70) | 5.58 (4.89) | 5.71 (4.53) | $p = .502$ |
| Baseline HPT<sub>h</sub> (°C) | 44.83 (3.55) | 44.54 (3.75) | 45.12 (3.33) | $p = .340$ |
| Baseline HPT<sub>o</sub> (°C) | 48.45 (1.49) | 48.22 (1.96) | 48.67 (0.74) | $p = .835$ |
| Post-MAST subjective stress levels (0-100) | 34.72 (31.53) | 58.27 (24.74) | 11.79 (17.29) | **$p < .001$** |
| Post-MAST subjective pain levels (0-100) | 34.74 (32.96) | 62.14 (21.84) | 8.01 (15.25) | **$p < .001$** |
| Post-MAST subjective unpleasantness levels (0-100) | 44.96 (34.56) | 72.63 (21.65) | 18.03 (20.65) | **$p < .001$** |
| Post-MAST positive affect (PANAS; 10-50) | 20.50 (4.25) | 22.01 (4.27) | 18.95 (3.61) | **$p < .001$** |
| Post-MAST negative affect (PANAS; 10-50) | 21.49 (4.11) | 22.15 (3.97) | 20.84 (4.17) | $p = .073$ |
| MAP reactivity ($\Delta$ MAP) | -0.01 (8.69) | 2.85 (7.46) | -2.79 (8.93) | **$p < .001$** |
| Pulse reactivity ($\Delta$ Pulse) | -2.53 (18.03) | -6.42 (17.62) | 1.26 (17.72) | **$p = .020$** |
| Alpha-amylase reactivity ($\Delta$ Alpha-amylase) | 14.37 (54.76) | 28.96 (54.69) | -0.02 (51.24) | **$p < .001$** |
| Cortisol reactivity ($\Delta$ Cortisol) | 1.91 (6.36) | 5.42 (6.76) | -1.50 (3.45) | **$p < .001$** |
| AUCi cortisol | 31.00 (175.08) | 107.91 (185.26) | -42.84 (127.56) | **$p < .001$** |
| Stress-induced changes in HPT<sub>h</sub> ($\Delta$ HPT<sub>h</sub>) | 0.47 (2.08) | 0.23 (2.08) | 0.71 (2.06) | $p = .075$ |
| Stress-induced changes in HPT<sub>o</sub> ($\Delta$ HPT<sub>o</sub>) | -0.07 (0.60) | -0.13 (0.73) | -0.01 (0.45) | $p = .406$ |

In the stress group: (a) 50 participants (68.5%) had an increase of ≥1.5 nmol/l due to stress induction and were classified as cortisol stress responders, (b) 22 participants (30.1%) were characterized by stress-induced hyperalgesia (SI-hyper), 16 participants (21.9%) by stress-induced hypoalgesia (SI-hypo), and 35 participants (48%) as stress-induced non-responder (SI-NR). These subgroups were used in exploratory analyses.

### 3.2 Stress manipulation checks

An overview of the stress manipulation checks can be found in **Figure 2**.

**Figure 2.**
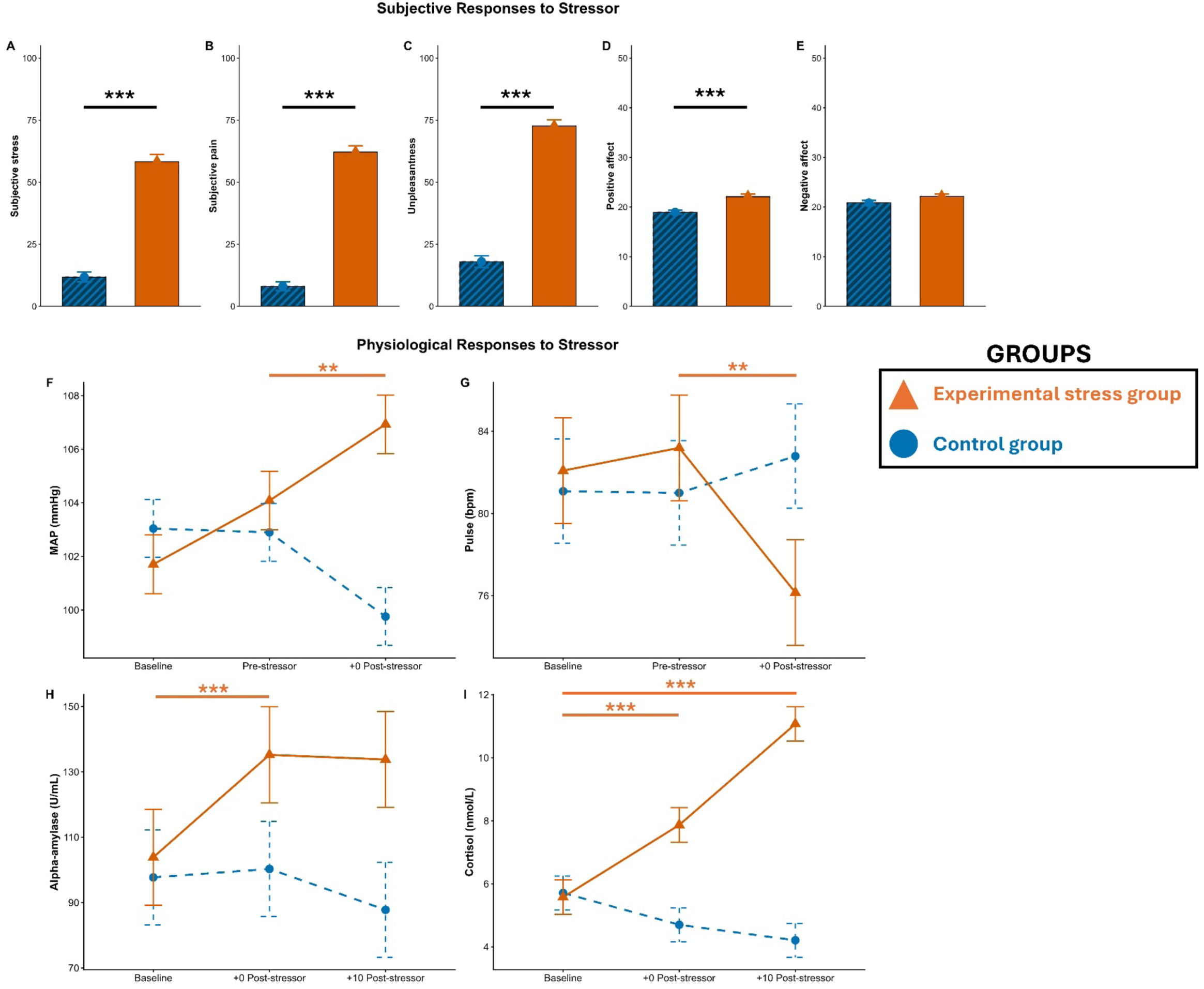
Overview of the stress manipulation checks. MAP: Mean Arterial Pressure, MAST: Maastricht Acute Stress Task, *: p < .05, **: p < .01, ***: p < .001. To facilitate interpretation, the figure depicts raw values and not log-transformed values (created with ggplot2 package in RStudio). VAS-scores for stress (**A**), pain (**B**), unpleasantness (**C**) and positive (**D**) and negative affect (**E**) using the PANAS were assessed after the MAST. MAP (**F**) and pulse (**G**) were measured at baseline, before introduction of the MAST and immediately after the MAST. Salivary alpha-amylase (**H**) and cortisol (**I**) values were collected at baseline, immediately after MAST, and 10 minutes after the MAST.

Post-MAST subjective stress, pain and unpleasantness were significantly higher in the stress group compared to the control group as indicated by Wilcoxon rank-sum test (*W* = 396, *W* = 213, *W* = 303 respectively, all *p* < .001), post-MAST positive affect was significantly higher in the stress group compared to the control group as shown by an independent samples *t*-test (*t* = -4.863, df = 142.61, *p* < .001), and post-MAST negative affect did not differ between groups (*W* = 227.5, *p* = .073).

For MAP, a significant interaction effect between time and group [*F*_2, 292_ = 20.51, *p* < .001, *η²ₚ* = 0.12] was found, while main effects of time [*F*_2, 292_ = 2.42, *p* = .091, *η²ₚ* = 0.02] and group [*F*_1, 146_ = 2.88, *p* = .092, *η²ₚ* = 0.02] were not significant. Post-hoc *t*-tests showed no baseline group differences (estimate = 1.335, SE = 1.460, df = 246, 95% CI [-1.540, 4.210], *t* = 0.916, *p* = .36), whereas directly after the stressor, the groups differed significantly (estimate = -6.578, SE = 1.460, df = 246, 95% CI [-9.450, -3.710], *t* = -4.514, *p* < .001). Only in the stress group, a significant increase in MAP over time occurred (estimate = -2.852, SE = 0.899, df = 292, 95% CI [-4.620, -1.083], *t* = -3.174, *p* = .002, *pHolm = .*003).

For pulse, the interaction effect between time and group became significant [*F*_2, 292_ = 5.58, *p* = .004, *η²ₚ* = 0.04], while no significant main effects of time [*F*_2, 292_ = 2.34, *p* = .098, *η²ₚ* = 0.02] and group [*F*_1, 146_ = 0.09, *p* = .77, *η²ₚ* < 0.01] were observed. Post-hoc *t*-tests showed no group differences at baseline (estimate = -0.016, SE = 0.038, df = 212, 95% CI [-0.091, 0.060], *t* = - 0.408, *p* = .68) or immediately after the stressor (estimate = 0.064, SE = 0.038, df = 212, 95% CI [-0.011, 0.140], *t* = 1.685, *p* = .094). Pulse values decreased significantly over time in the stress group (estimate = 0.069, SE = 0.020, df = 292, 95% CI [0.030, 0.101], *t* = 3.442, *p* < .001, *pHolm = .*002), whereas they remained stable in the control group.

For alpha-amylase, a significant interaction effect between time and group [*F*_1, 142.19_ = 17.21, *p* < .001, *η²ₚ* = 0.11] as well as a significant main effect of time [*F*_1, 142.19_ = 4.09, *p* = .045, *η²ₚ* = 0.03] was observed, while no significant main effect of group [*F*_1, 143.90_ = 2.31, *p* = .13, *η²ₚ* = 0.02] was present. Post-hoc *t*-tests showed no baseline group differences (estimate = -0.054, SE = 0.161, df = 167, 95% CI [-0.372, 0.264], *t* = -0.337, *p* = .74), but a significant group difference evolved immediately post-stressor (estimate = -0.417, SE = 0.161, df = 168, 95% CI [-0.735, -0.094], *t* = -2.584, *p* = .011). In the stress group, a significant alpha-amylase increase from baseline to immediately post-stressor (estimate = -0.270, SE = 0.062, df = 143, 95% CI [-0.392, -0.147], *t* = -4.349, *p* < .001) was detected. This was not the case in the control group.

For cortisol, significant main effects of group [*F*_1, 137.84_ = 18.15, *p* < .001, *η²ₚ* = 0.12], time [*F*_2, 274.95_ = 23.34, *p* < .001, *η²ₚ* = 0.15], and interaction between time and group [*F*_2, 274.95_ = 77.84, *p* < .001, *η²ₚ* = 0.36] were observed. Simple effects showed no significant group differences at baseline (estimate = 0.122, SE = 0.113, df = 205, 95% CI [-0.099, 0.344], *t* = 1.086, *p* = .28), whereas a significant difference was present immediately after stressor (estimate = -0.477, SE = 0.113, df = 206, 95% CI [-0.700, -0.255], *t* = -4.230, *p* < .001) and 10 minutes post-stressor (estimate = -0.940, SE = 0.113, df = 205, 95% CI [-1.162, -0.718], *t* = -8.345, *p* < .001). In the stress group, a significant increase from baseline to immediately post-stressor (estimate = -0.457, SE = 0.062, df = 275, 95% CI [-0.605, -0.309], *t* = -7.427, both *p* and *pHolm* < .001), and from baseline to 10 minutes post-stressor (estimate = -0.822, SE = 0.061, df = 275, 95% CI [-0.970, -0.675], *t* = -13.434, both *p* and *pHolm* < .001) evolved. This was also the case for the control group, but in the other direction (i.e., decrease from baseline to immediately after stressor: estimate = 0.142, SE = 0.060, df = 275, 95% CI [-0.0009, 0.286], *t* = 2.393, *p* = .017, *pHolm* = .035; and decrease from baseline to 10 minutes post-stressor: estimate = 0.239, SE = 0.060, df = 275, 95% CI [0.096, 0.383], *t* = 4.031, both *p* and *pHolm* < .001).

### 3.3 Associations between trait pain-related distress and stress reactivity (O1)

To test whether individual differences in trait pain-related distress were associated with variations in stress reactivity, correlation analyses were conducted. Focusing on the stress group only, the score of TSK was negatively correlated with the change in alpha-amylase (*r* = -.276, *p* = .019, uncorrected; *p-FDR* = .23). No other significant correlations were found (all *p* > .17, uncorrected) and no associations survived the FDR-correction. When zooming in on cortisol stress responders only, again no significant associations were found (all *p* > .23, uncorrected). A complete overview of the results can be found in **Supplementary Table 1**.

Also, no significant differences in pain-related distress were observed between the three subgroups of pain (i.e., SI-hypo, SI-hyper, SI-NR, all *p* > .10, uncorrected). Finally, and only for completeness, we also explored potential associations between trait pain-related distress and baseline physiological stress markers. These are listed in the **Supplementary Results** and **Supplementary Table 2**.

### 3.4 Pain sensitivity and stress reactivity

#### 3.4.1 Associations between (stress-induced changes in) pain sensitivity and stress reactivity (O2)

To test whether individual differences in baseline pain sensitivity and changes in pain sensitivity after acute stress were associated with variations in stress reactivity, correlation analyses were conducted. Respective results revealed that stress reactivity did not correlate with (changes in) pain thresholds and tolerances (all *p* > .07, uncorrected). Exploratory analyses in cortisol responders only did not yield any significant findings (all *p* > .18, uncorrected). A graphical representation of how pain thresholds change over time for each person can be found in **Figure 3** and an overview of all results is provided in **Supplementary Table 3**.

**Figure 3.**
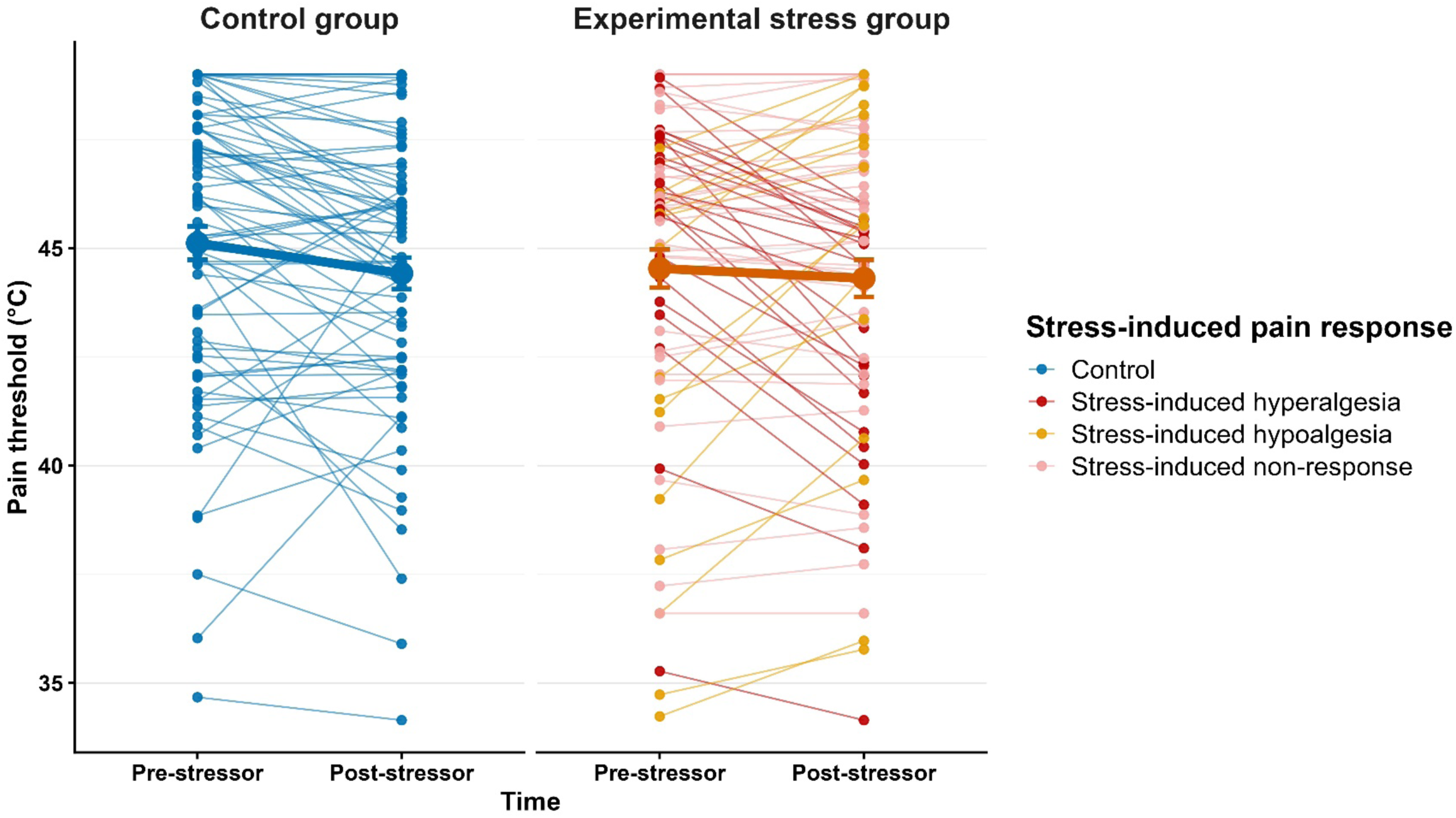
Individual differences in pain threshold before and after acute stress induction. Spaghetti plots indicating the variability in the individual pain thresholds in response to the stressor (created with ggplot2 package in RStudio).

Also, no significant differences in stress reactivity were observed between the three subgroups of pain (i.e., SI-hypo, SI-hyper, SI-NR, all *p* > .17, uncorrected) Finally, explorative results on the associations between pain thresholds and tolerances and baseline stress markers are reported in the **Supplementary Results** and **Supplementary Table 4**.

#### 3.4.2 Group-level changes in pain sensitivity over time

For pain thresholds, no interaction effect between time and group [*F*_1, 146_ = 1.840, *p* = .18, *η²ₚ* = 0.01], and no main effect of group [*F*_1, 146_ = 0.409, *p* = .52, *η²ₚ* < 0.001] were observed, but the main effect of time [*F*_1, 146_ = 7.12, *p* = .008, *η²ₚ* = 0.05] was significant. Exploratively, only in the control group, pain thresholds decreased significantly from baseline to post-stressor (estimate = 0.697, SE = 0.243, df = 146, 95% CI [0.216, 1.177], *t* = 2.866, *p* = 0.048). No significant group differences at baseline or after the stressor were detected (all *p* > .31).

For pain tolerance, no significant interaction effect between time and group [*F*_1, 146_ = 3.641, *p* = .058, *η²ₚ* = 0.02] and no significant main effect of group [*F*_1, 146_ = 1.909, *p* = .17, *η²ₚ* = 0.01] was present, but the main effect of time [*F*_1, D146_ = 4.56, *p* = .034, *η²ₚ* = 0.03] reached significance. Explorative analyses revealed that only in the stress group, pain tolerances increased significantly (estimate = -0.178, SE = 0.063, df = 146, 95% CI [-0.302, -0.054], *t* = - 2.840, *p* = .005). No significant group differences existed at baseline or post-stressor (all *p* > .07).

### 3.5 Trait pain-related distress as a moderator of the relationship between stress reactivity and stress-induced changes in pain sensitivity (O3)

To examine whether individual differences in trait pain-related distress moderate the relationship between stress reactivity and stress-induced changes in pain sensitivity, moderation analyses were conducted (in the stress group only). Moderation analyses revealed that in individuals with higher trait pain catastrophizing and higher trait fear of pain, larger alpha-amylase increases were associated with larger post-stressor increases in pain threshold. More specifically, the change in alpha-amylase X FPQ interaction term (*B* = –0.001, SE < 0.001, 95% CI [–0.002, –0.000], *t*(68) = –2.58, *p* = .012, *p-FDR* = .072*, β* = –.29, Δ*R*² = .09, *R*² = .11), and the change in alpha-amylase X PCS interaction term (*B* = –0.002, 95% bootstrapped CI [–0.004, -0.000], *p* = .017, *p-FDR* = .072, β = –.40, Δ*R*² = .09, *R*² = .10) were significant at an uncorrected *p*-level, but did not survive FDR-correction. Simple slopes of these relationships can be retrieved in **Figure 4**. No other interaction effects were significant. Exploratory analyses in cortisol responders only did not change results. A complete overview of all moderation analyses is provided in **Supplementary Table 5.**

**Figure 4.**
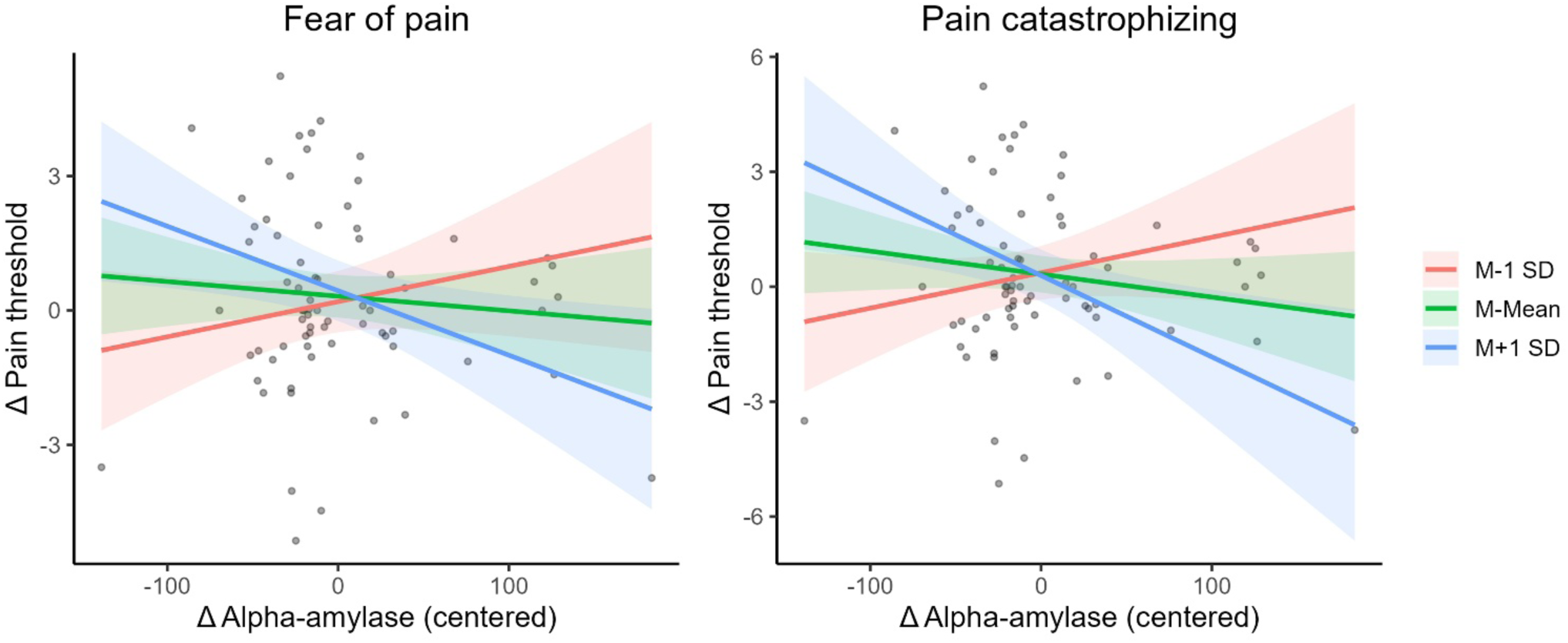
Fear of pain and pain catastrophizing moderate the association between alpha-amylase reactivity and pain threshold change. M = Mean, SD = Standard deviation, Δ = change in. Simple slopes of the relationship between stress-induced alpha-amylase reactivity and change in pain threshold at low (−1 SD), mean, and high (+1 SD) levels of trait fear of pain (FPQ; left) and trait pain catastrophizing (PCS; right). At higher FPQ or PCS, higher alpha-amylase reactivity predicted larger increases in pain threshold (hypoalgesia), whereas no significant associations were observed at lower levels of these traits. Δ Pain threshold is expressed as Pain threshold_pre-stressor_ – Pain threshold_post-stressor_.

## 4 Discussion

The aim of this study was (a) to investigate whether individual differences in trait pain-related distress (i.e., pain catastrophizing, fear of movement, fear of pain) relate to physiological stress reactivity (i.e., operationalized as pre- to post-MAST changes in MAP, pulse, alpha-amylase and cortisol), (b) to determine whether stress reactivity is related to stress-induced changes in pain sensitivity (i.e., pre- to post-MAST changes in heat pain threshold and tolerance levels), and (c) to test whether trait pain-related distress moderates the relationship between stress reactivity and stress-induced changes in pain sensitivity. Across these objectives, the results reveal a complex and highly heterogeneous pattern of associations, highlighting that the stress-pain relationship cannot be captured by uniform group-level analyses.

More specifically, our results suggest that (a) there are no strong associations between trait pain-related distress and stress reactivity, as no associations survived multiple comparison correction, (b) there is high individual variability in how acute stress alters pain sensitivity, and (c) there is preliminary evidence suggesting that among individuals with a higher tendency to catastrophize pain or higher fear of pain, larger alpha-amylase increases may be associated with larger post-stressor increases in pain threshold, which is indicative of lower pain sensitivity (i.e., hypoalgesia). However, these results should be interpreted cautiously and require further study, as these findings did not survive correction for multiple comparisons.

### Effectiveness of the stress manipulation

First, manipulation checks of the stressor should be evaluated. The observed reactivity of blood pressure, alpha-amylase and cortisol to the MAST in the stress group is consistent with previous research using the MAST (Shilton et al., 2017; Smeets et al., 2012; Timmers et al., 2018). Also in line with earlier findings, no significant increase in pulse was found (Shilton et al., 2017; Timmers et al., 2018). Participants in the stress group reported higher levels of self-perceived stress, pain, and unpleasantness compared to the control group, confirming that the MAST successfully induced acute self-reported stress. Positive affect was significantly higher in the stress group, potentially reflecting relief once the task was completed. Overall, these findings indicate that the MAST elicited distinct SAM and HPA axes’ activation, providing a solid basis for examining individual variation in downstream pain modulation.

### Trait pain-related distress and stress reactivity

Our first preregistered objective was to investigate whether trait pain-related distress is associated with stress reactivity. Contrary to our preregistered hypothesis that individuals reporting more trait pain-related distress would exhibit greater physiological reactivity to experimentally induced stress, our results indicate no significant associations between pain-related distress and stress reactivity. At an uncorrected level, our findings suggest that individuals with a lower tendency to fear movement reacted with larger increases in alpha-amylase to acute stress, but this finding did not survive FDR-correction and hence should be interpreted with caution. Future studies should investigate whether there may indeed be different patterns between the more cognitive-emotional traits (fear of pain, catastrophizing) and more behaviourally oriented pain-related distress traits (kinesiophobia), with potentially only the behavioural action tendencies being associated with the latter playing a role in shaping acute physiological stress responses. Specifically, individuals with high fear of movement may adopt more inhibited or avoidant behavioural tendencies in novel or challenging situations (Meulders, 2019), which could dampen sympathetic reactivity (Evans et al., 2019). Future research with behavioural engagement measures could test this possibility more explicitly.

### Stress reactivity and changes in pain sensitivity

Secondly, we investigated the effect of an acute stressor on pain sensitivity and how these changes in pain sensitivity are related to physiological stress reactivity. We detected large individual differences in changes of pain thresholds and tolerances following the stress manipulation, without a group-level effect. Moreover, the strength of physiological stress responses did not predict these changes, which did not align with our hypothesis that greater stress responses would be associated with greater stress-induced changes in pain sensitivity. Although studies by Timmers et al. (2018) and Al’Absi and Petersen (2003) support the involvement of stress system reactivity of the SAM and HPA axes in stress-induced changes in pain sensitivity, more specifically in pain reduction, not all research supports this directionality. In fact, findings are mixed, with some studies indeed demonstrating stress-induced hyperalgesia instead (Crettaz et al., 2013; Reinhardt et al., 2013). Our findings are in line with other studies that also did not observe any significant changes in pain sensitivity following acute stress (Geva et al., 2014). Even within a single study, all three patterns (i.e., stress-induced hyperalgesia, hypoalgesia, and no effect on pain thresholds) have been identified (Bement et al., 2010; Gera et al., 2025), including in ours, highlighting substantial individual variability in stress–pain interactions. Importantly, these studies did not detect a significant overall effect (averaged across participants) of the acute stressor on pain thresholds, which is consistent with our study. Taken together, our findings support the growing notion that stress-related pain modulation reflects a modular and functionally dissociable system in which SAM activation, HPA axis activity, and endogenous pain modulation do not necessarily operate in synchrony. Another potential explanation for the null finding is that the substantial individual heterogeneity in stress-induced effects on pain is better explained by other factors than activation of the SAM and HPA axes that were not captured in our study, including for instance appraisals or mindsets related to stress. The importance of individual differences in stress-induced pain responses is clear though, and the underlying mechanisms merit further investigation.

### Moderating role of trait pain-related distress

Our study also examined the moderating role of trait pain-related distress in the association between stress reactivity and stress-induced changes in pain sensitivity. Our study presents preliminary evidence suggesting that among individuals with higher fear of pain and pain catastrophizing, stronger alpha-amylase reactivity (i.e., sympathetic reactivity) predicted greater hypoalgesia, whereas no such relationship was observed among individuals with lower distress. This finding does not align with our hypothesis that greater stress responses would be associated with hypoalgesia in individuals with low pain-related distress. Instead, it suggests that for healthy volunteers, experiencing pain-related distress may be adaptive to some extent, as for those individuals, activating the sympathetic nervous system might have protective effects on pain, while for those with lower pain-related distress, it does not. Whether this effect also applies to clinical samples that experience substantially higher levels of pain-related distress remains to be tested. In this regard, previous studies revealed that higher arousal is associated with higher activation of the descending pain-inhibitory pathways (Rhud and Meagher, 2001), and that SAM reactivity contributes to stress-induced reductions in pain sensitivity (Al’Absi and Petersen, 2003). Our findings are in line with these studies in also supporting the proposed role for the SAM system, but not for the HPA axis, thus that we conclude that the patterns are more nuanced. And even though our findings are seemingly in contrast with findings by Timmers et al. (2018), who reported that the HPA axis, rather than the SAM axis, underlies stress-induced hypoalgesia, it should be noted that in our study pain thresholds and tolerances were reassessed directly after the end of the stressor (i.e., within the SAM axis response window), when cortisol levels have likely not peaked yet. It is also important to note that it has been shown that the nature of the stressor can differentially influence SAM and HPA axes reactivity (Brunyé et al., 2025). We therefore recommend future research to explore the dependency of the stress axes’ reactivity on the type of the stressor in more detail.

### Individual variability as a central feature of the stress-pain system

Across all objectives, one consistent observation was that the magnitude and direction of both the stress responses and pain modulation varied widely between individuals. Moreover, we detected evidence suggesting that psychological traits may moderate or even shape the impact of physiological stress reactivity on pain outcomes, rather than uniformly affecting stress physiology. Other researchers have also demonstrated that group comparisons can obscure important heterogeneity within the population (Bement et al., 2010; Gera et al., 2025; Hilger and Hewig, 2022; Reinhardt et al., 2013), thus that accounting for such individual differences, particularly in traits related to stress and pain, should become essential in future research on stress, pain, as well as on the link between stress reactivity and stress-induced changes in pain sensitivity.

Finally, such variability also likely contributes to the inconsistencies in the stress-pain literature, where studies alternately report hyperalgesia, hypoalgesia, or null findings after acute stress. The present results, therefore, underline calls for more person-centred analytic approaches, such as idiographic modelling, dynamic systems perspectives, and the study of latent responder subtypes.

### Strength and limitations

The strength of our study includes strict eligibility criteria and a highly standardized design that reduced variability. Furthermore, a validated stressor was used, and manipulation checks confirmed its effectiveness. However, some limitations need to be addressed. First, the thermal stimulator introduced a practical limit for assessing pain tolerance (maximum temperature: 49 °C), which constrains the interpretation of stress-related changes in pain tolerance. Second, we cannot rule out that repeated pain assessment within the same day might have introduced habituation or carry-over effects, although such effects could not explain the variability we observe. Third, while our focus was on physiological stress markers that were assessed repeatedly throughout the experiment, subjective stress was measured only once following the stressor (as a manipulation check), which restricts the ability to capture temporal dynamics of the subjective stress responses and thus to examine its role more in detail. Fourth, the homogeneity of the sample limits generalizability beyond those assigned male at birth, even though sex differences in stress reactivity (Liu et al., 2017) and pain modulation (Keogh, 2022) have been documented. Thus, we cannot expect that the observed observations and null results translate to a female sample, where for example hormonal influences could modify the effects (Gu et al., 2022), or to clinical populations, where dysregulations may amplify or obscure the stress-pain associations. Lastly, we acknowledge that the psychometric properties of the FPQ and TSK raise potentials concerns, and that the PCS captures both trait and state characteristics (although more trait contribution in a younger sample; Dumenci et al., 2020). Nonetheless, these instruments are widely used and were appropriate for our study, given their availability in German and suitability for a non-clinical population.

## 5 Conclusion

Taken together, this study demonstrates that the dynamics between trait pain-related distress, stress physiology, and stress-induced pain sensitivity follow idiosyncratic patterns. We observed no significant associations between pain-related distress and physiological stress reactivity, and stress reactivity was not related to stress-induced changes in pain sensitivity. However, our study provides preliminary evidence that among individuals with a higher tendency to catastrophize and/or fear pain, greater sympathetic reactivity to an acute stressor is associated with larger stress-induced increases in pain thresholds (indicating reduced pain sensitivity or hypoalgesia). While these findings did not survive correction for multiple testing and should therefore be interpreted with caution, our study suggests that individual differences may shape stress-pain interactions. This also implies that group-level analyses that aim to explain stress-pain interactions by physiological responses alone may fall short or detect inconsistent fragile patterns, lacking replication. We propose that future research on stress-pain interactions should put an explicit focus on the individual level to unravel potential moderating factors (e.g., personality traits, past experiences), which could support the translation of findings on stress-pain interactions to chronic pain populations, with the ultimate goal of developing novel tailored prevention and treatment programs.

## Declaration of generative AI and AI-assisted technologies in the manuscript preparation process

During the preparation of this work the author(s) used ChatGPT to assist in developing the code in the script in R. After using this tool, the author(s) reviewed and edited the content as needed and take full responsibility for the content of the published article.

## Author contributions

Conceptualization: Inge Timmers, Joren Vyverman, Kirsten Hilger, Stefanie H Meeuwis, Tom Smeets.

Data curation: Kirsten Hilger

Formal analysis: Joren Vyverman

Funding acquisition: Kirsten Hilger

Investigation: Kirsten Hilger

Methodology: Kirsten Hilger

Project administration: Kirsten Hilger

Resources: Kirsten Hilger

Supervision: Kirsten Hilger

Validation: Joren Vyverman, Kirsten Hilger

Visualization: Joren Vyverman

Writing – original draft: Joren Vyverman

Writing – Review & editing: Inge Timmers, Joren Vyverman, Kirsten Hilger, Stefanie H Meeuwis, Tom Smeets.

All authors contributed to and have approved the final version of the manuscript.

## Supplementary material

Supplementary materials are available.

## References

Aiken, L.S., West, S.G., Reno, R.R., 1991. Multiple regression: Testing and interpreting interactions. sage.

Al’Absi, M., Petersen, K.L., 2003. Blood pressure but not cortisol mediates stress effects on subsequent pain perception in healthy men and women. Pain 106, 285–295.

Arendt-Nielsen, L., Yarnitsky, D., 2009. Experimental and clinical applications of quantitative sensory testing applied to skin, muscles and viscera. The journal of pain 10, 556–572.

Baum, C., Schneider, R., Keogh, E., Lautenbacher, S., 2013. Different stages in attentional processing of facial expressions of pain: a dot-probe task modification. The Journal of Pain 14, 223–232.

Bement, M.H., Weyer, A., Keller, M., Harkins, A.L., Hunter, S.K., 2010. Anxiety and stress can predict pain perception following a cognitive stress. Physiology & behavior 101, 87–92.

Besson, J., 1999. The neurobiology of pain. The Lancet 353, 1610–1615.

Breyer, B., Bluemke, M., 2016. Deutsche version der positive and negative affect schedule PANAS (GESIS panel).

Brunyé, T.T., Goring, S.A., Navarro, E., Hart-Pomerantz, H., Grekin, S., McKinlay, A.M., Plessow, F., 2025. Identifying the most effective acute stress induction methods for producing SAM-and HPA-related physiological responses: a meta-analysis. Anxiety, Stress, & Coping 38, 263–285.

Cardinali, D.P., 2018. Autonomic nervous system. Springer International Publishing 10, 978–973.

Crettaz, B., Marziniak, M., Willeke, P., Young, P., Hellhammer, D., Stumpf, A., Burgmer, M., 2013. Stress-induced allodynia–evidence of increased pain sensitivity in healthy humans and patients with chronic pain after experimentally induced psychosocial stress. PloS one 8, e69460.

Crombez, G., Eccleston, C., Van Damme, S., Vlaeyen, J.W., Karoly, P., 2012. Fear-avoidance model of chronic pain: the next generation. The Clinical journal of pain 28, 475–483.

Dumenci, L., Kroenke, K., Keefe, F.J., Ang, D.C., Slover, J., Perera, R.A., Riddle, D.L., 2020. Disentangling trait versus state characteristics of the Pain Catastrophizing Scale and the PHQ-8 Depression Scale. European Journal of Pain 24, 1624–1634.

Evans, T.C., Rodriguez, A.M., Britton, J.C., 2019. Sympathetic and self-reported threat reactivity in social anxiety: Modulation by threat certainty and avoidance behavior. Journal of Psychopathology and Behavioral Assessment 41, 627–638.

Faul, F., Erdfelder, E., Buchner, A., Lang, A.-G., 2009. Statistical power analyses using G* Power 3.1: Tests for correlation and regression analyses. Behavior research methods 41, 1149–1160.

Gebhart, G., 2004. Descending modulation of pain. Neuroscience & Biobehavioral Reviews 27, 729–737.

Gera, O., Ginzburg, K., Gur, N., Defrin, R., 2025. Effects of acute stress exposure on pain sensitivity: the role of individual stress responsiveness and orientation to pain and stress. Pain 166, e388–e396.

Geva, N., Defrin, R., 2018. Opposite effects of stress on pain modulation depend on the magnitude of individual stress response. The Journal of Pain 19, 360–371.

Geva, N., Pruessner, J., Defrin, R., 2014. Acute psychosocial stress reduces pain modulation capabilities in healthy men. Pain® 155, 2418–2425.

Hilger, K., Häge, A.-S., Zedler, C., Jost, M., Pauli, P., 2023. Virtual reality to understand pain-associated approach behaviour: a proof-of-concept study. Scientific Reports 13, 13799.

Hilger, K., Hewig, J., 2022. Individual differences in the focus: Understanding variations in pain-related fear and avoidance behavior from the perspective of personality science. Pain 163, e151–e152.

Houben, R.M., Leeuw, M., Vlaeyen, J.W., Goubert, L., Picavet, H.S.J., 2005. Fear of movement/injury in the general population: factor structure and psychometric properties of an adapted version of the Tampa Scale for Kinesiophobia. Journal of behavioral medicine 28, 415–424.

Keogh, E., 2022. Sex and gender differences in pain: past, present, and future. Pain 163, S108–S116.

Kirschbaum, C., Pirke, K.-M., Hellhammer, D.H., 1993. The ‘Trier Social Stress Test’–a tool for investigating psychobiological stress responses in a laboratory setting. Neuropsychobiology 28, 76–81.

Kudielka, B.M., Hellhammer, D.H., Wüst, S., 2009. Why do we respond so differently? Reviewing determinants of human salivary cortisol responses to challenge. Psychoneuroendocrinology 34, 2–18.

Liu, J.J., Ein, N., Peck, K., Huang, V., Pruessner, J.C., Vickers, K., 2017. Sex differences in salivary cortisol reactivity to the Trier Social Stress Test (TSST): A meta-analysis. Psychoneuroendocrinology 82, 26–37.

McNeil, D.W., Rainwater, A.J., 1998. Development of the fear of pain questionnaire-III. Journal of behavioral medicine 21, 389–410.

Meaney, E., Alva, F., Moguel, R., Meaney, A., Alva, J., Webel, R., 2000. Formula and nomogram for the sphygmomanometric calculation of the mean arterial pressure. Heart 84, 64–64.

Meulders, A., 2019. From fear of movement-related pain and avoidance to chronic pain disability: a state-of-the-art review. Current Opinion in Behavioral Sciences 26, 130–136.

Meyer, K., Sprott, H., Mannion, A.F., 2008. Cross-cultural adaptation, reliability, and validity of the German version of the Pain Catastrophizing Scale. Journal of psychosomatic research 64, 469–478.

Miller, R., Plessow, F., Kirschbaum, C., Stalder, T., 2013. Classification criteria for distinguishing cortisol responders from nonresponders to psychosocial stress: evaluation of salivary cortisol pulse detection in panel designs. Psychosomatic medicine 75, 832–840.

Orloff, S., 1979. Pain tolerance threshold and pain perception threshold: clinical and experimental studies. Clinics in Rheumatic Diseases 5, 755–773.

Pruessner, J.C., Kirschbaum, C., Meinlschmid, G., Hellhammer, D.H., 2003. Two formulas for computation of the area under the curve represent measures of total hormone concentration versus time-dependent change. Psychoneuroendocrinology 28, 916–931.

R Core Team, 2025. R: A Language and Environment for Statistical Computing. R Foundation for Statistical Computing, Vienna, Austria.

Raja, S.N., Carr, D.B., Cohen, M., Finnerup, N.B., Flor, H., Gibson, S., Keefe, F.J., Mogil, J.S., Ringkamp, M., Sluka, K.A., 2020. The revised International Association for the Study of Pain definition of pain: concepts, challenges, and compromises. Pain 161, 1976–1982.

Reinhardt, T., Kleindienst, N., Treede, R.-D., Bohus, M., Schmahl, C., 2013. Individual modulation of pain sensitivity under stress. Pain Medicine 14, 676–685.

Rhud, J.L., Meagher, M.W., 2001. Noise stress and human pain thresholds: divergent effects in men and women. The Journal of Pain 2, 57–64.

Rohleder, N., Nater, U.M., Wolf, J.M., Ehlert, U., Kirschbaum, C., 2004. Psychosocial stress-induced activation of salivary alpha-amylase: an indicator of sympathetic activity? Annals of the New York Academy of Sciences 1032, 258–263.

Rohleder, N., Wolf, J.M., Maldonado, E.F., Kirschbaum, C., 2006. The psychosocial stress-induced increase in salivary alpha-amylase is independent of saliva flow rate. Psychophysiology 43, 645–652.

Rosenblum, S., Rab, S.L., Admon, R., 2025. Dynamics in physiological acute stress response trajectories: uncovering latent variability. BMC psychiatry 25, 361.

Schlotz, W., Hammerfald, K., Ehlert, U., Gaab, J., 2011. Individual differences in the cortisol response to stress in young healthy men: Testing the roles of perceived stress reactivity and threat appraisal using multiphase latent growth curve modeling. Biological psychology 87, 257–264.

Shilton, A.L., Laycock, R., Crewther, S.G., 2017. The Maastricht Acute Stress Test (MAST): Physiological and subjective responses in anticipation, and post-stress. Frontiers in psychology 8, 567.

Smeets, T., Cornelisse, S., Quaedflieg, C.W., Meyer, T., Jelicic, M., Merckelbach, H., 2012. Introducing the Maastricht Acute Stress Test (MAST): a quick and non-invasive approach to elicit robust autonomic and glucocorticoid stress responses. Psychoneuroendocrinology 37, 1998–2008.

Smeets, T., Van Ruitenbeek, P., Hartogsveld, B., Quaedflieg, C.W., 2019. Stress-induced reliance on habitual behavior is moderated by cortisol reactivity. Brain and Cognition 133, 60–71.

Soliemanifar, O., Soleymanifar, A., Afrisham, R., 2018. Relationship between personality and biological reactivity to stress: a review. Psychiatry investigation 15, 1100.

Sullivan, M.J., Bishop, S.R., Pivik, J., 1995. The pain catastrophizing scale: development and validation. Psychological assessment 7, 524.

Timmers, I., Kaas, A.L., Quaedflieg, C.W., Biggs, E.E., Smeets, T., de Jong, J.R., 2018. Fear of pain and cortisol reactivity predict the strength of stress-induced hypoalgesia. European Journal of Pain 22, 1291–1303.

Ulrich-Lai, Y.M., Herman, J.P., 2009. Neural regulation of endocrine and autonomic stress responses. Nature reviews neuroscience 10, 397–409.

Von Elm, E., Altman, D.G., Egger, M., Pocock, S.J., Gøtzsche, P.C., Vandenbroucke, J.P., 2007. The Strengthening the Reporting of Observational Studies in Epidemiology (STROBE) statement: guidelines for reporting observational studies. The lancet 370, 1453–1457.

Watson, D., Clark, L.A., Tellegen, A., 1988. Development and validation of brief measures of positive and negative affect: the PANAS scales. Journal of personality and social psychology 54, 1063.

