## Supplementary for "Individual Dynamics in Stress-Related Pain Responses"

### Supplementary Results

#### Trait pain-related distress and the baseline markers of stress

To test whether individual differences in trait pain-related distress were associated with baseline stress markers across all participants, correlation analyses were conducted. A summary of the results can be found in **Supplementary Table 2**.

TSK scores were positively associated with baseline alpha-amylase values ($r$ = .167, $p$ = .043, uncorrected; *p-FDR* = .257), and FPQ scores were negatively associated with baseline MAP values ($r$ = -.177, $p$ = .032, uncorrected; *p-FDR* = .257). No other significant correlations were found (all $p$ > .16, uncorrected) and no associations survived the FDR-correction.

#### Baseline pain sensitivity and the baseline markers of stress

To test whether baseline pain sensitivity was associated with baseline stress markers across all participants, correlation analyses were conducted. A summary of the results can be found in **Supplementary Table 4.**

Baseline pain thresholds were significantly associated with baseline MAP values ($r$ = .179, $p$ = .030, uncorrected; *p-FDR* = .198). No other significant correlations were found (all $p$ > .06, uncorrected) and no associations survived the FDR-correction.

**Supplementary Table 1. Correlations between trait pain-related distress and stress reactivity.** *FPQ = Fear of Pain Questionnaire, MAP = Mean Arterial Pressure, NA = not applicable, p(-FDR) = Probability value (adjusted for multiple comparisons using False Discovery Rate correction), PCS = Pain Catastrophizing Scale, r = Correlation Coefficient, TSK = Tampa Scale of Kinesiophobia.*

| Variable 1 | Variable 2 | *r* | *p-value* | *p-FDR* |
|---|---|---|---|---|
| PCS sum score | Δ MAP | .143 | .227 | .501 |
| TSK sum score | Δ MAP | .070 | .554 | .650 |
| FPQ sum score | Δ MAP | .063 | .596 | .650 |
| PCS sum score | Δ Pulse | .136 | .251 | .501 |
| TSK sum score | Δ Pulse | .141 | .235 | .501 |
| FPQ sum score | Δ Pulse | .147 | .215 | .501 |
| PCS sum score | Δ Alpha-amylase | .163 | .171 | .501 |
| TSK sum score | Δ Alpha-amylase | -.276 | **.019** | .226 |
| FPQ sum score | Δ Alpha-amylase | .036 | .764 | .764 |
| PCS sum score | Δ Cortisol | -.106 | .389 | .650 |
| TSK sum score | Δ Cortisol | -.082 | .509 | .650 |
| FPQ sum score | Δ Cortisol | -.083 | .502 | .650 |
| cortisol stress responders only | | | | |
| PCS sum score | Δ Cortisol | -.174 | .228 | NA |
| TSK sum score | Δ Cortisol | .003 | .985 | NA |
| FPQ sum score | Δ Cortisol | -.062 | .667 | NA |

**Supplementary Table 2. Correlations between trait pain-related distress and baseline stress markers of the entire sample.** *FPQ = Fear of Pain Questionnaire, MAP = Mean Arterial Pressure, p(-FDR) = Probability value (adjusted for multiple comparisons using False Discovery Rate correction), PCS = Pain Catastrophizing Scale, r = Correlation Coefficient, TSK = Tampa Scale of Kinesiophobia.*

| Variable 1 | Variable 2 | *r* | *p-value* | $p_{FDR}$ |
|---|---|---|---|---|
| PCS sum score | Baseline Cortisol | -.009 | .920 | .984 |
| TSK sum score | Baseline Cortisol | -.070 | .422 | .844 |
| FPQ sum score | Baseline Cortisol | -.002 | .984 | .984 |
| PCS sum score | Baseline Alpha-amylase | -.100 | .23 | .689 |
| TSK sum score | Baseline Alpha-amylase | .167 | **.043** | .257 |
| FPQ sum score | Baseline Alpha-amylase | .071 | .392 | .844 |
| PCS sum score | Baseline Pulse | .035 | .672 | .925 |
| TSK sum score | Baseline Pulse | -.008 | .927 | .984 |
| FPQ sum score | Baseline Pulse | .048 | .559 | .925 |
| PCS sum score | Baseline MAP | .0330 | .694 | .925 |
| TSK sum score | Baseline MAP | -.116 | .161 | .645 |
| FPQ sum score | Baseline MAP | -.177 | **.032** | .257 |

**Supplementary Table 3. Correlations between (changes in) pain sensitivity and stress reactivity.** *MAP = Mean Arterial Pressure, NA = not applicable, p(-FDR) = Probability value (adjusted for multiple comparisons using False Discovery Rate correction), r = Correlation Coefficient*

| Variable 1 | Variable 2 | *r* | *p-value* | *p-FDR* |
|---|---|---|---|---|
| *Changes in stress markers x pain sensitivity in the stress group* | | | | |
| Δ MAP | Pain Threshold | -.031 | .796 | .938 |
| Δ MAP | Pain Tolerance | .008 | .944 | .944 |
| Δ Pulse | Pain Threshold | -.216 | .067 | .533 |
| Δ Pulse | Pain Tolerance | .039 | .744 | .938 |
| Δ Alpha-amylase | Pain Threshold | .027 | .821 | .938 |
| Δ Alpha-amylase | Pain Tolerance | .096 | .423 | .938 |
| Δ Cortisol | Pain Threshold | .075 | .544 | .938 |
| Δ Cortisol | Pain Tolerance | .058 | .639 | .938 |
| cortisol stress responders only | | | | |
| Δ Cortisol | Pain Threshold | .171 | .234 | N.A. |
| Δ Cortisol | Pain Tolerance | -.019 | .897 | N.A. |
| *Changes in stress markers x changes in pain sensitivity in the stress group* | | | | |
| Δ MAP | Δ Pain Threshold | -.208 | .077 | .617 |
| Δ MAP | Δ Pain Tolerance | -.113 | .343 | .685 |
| Δ Pulse | Δ Pain Threshold | -.008 | .948 | .971 |
| Δ Pulse | Δ Pain Tolerance | .123 | .301 | .685 |
| Δ Alpha-amylase | Δ Pain Threshold | -.055 | .646 | .971 |
| Δ Alpha-amylase | Δ Pain Tolerance | .168 | .158 | .630 |
| Δ Cortisol | Δ Pain Threshold | .006 | .959 | .971 |
| Δ Cortisol | Δ Pain Tolerance | -.005 | .971 | .971 |
| cortisol stress responders only | | | | |
| Δ Cortisol | Δ Pain Threshold | .085 | .559 | NA |
| Δ Cortisol | Δ Pain Tolerance | -.126 | .385 | NA |

**Supplementary Table 4. Correlations between baseline pain sensitivity and baseline stress markers across the entire sample.** *MAP = Mean Arterial Pressure, p(-FDR) = Probability value (adjusted for multiple comparisons using False Discovery Rate correction), r = Correlation Coefficient*

| **Variable 1** | **Variable 2** | ***r*** | ***p-value*** | $p_{FDR}$ |
|---|---|---|---|---|
| Baseline Cortisol | Pain Threshold | .077 | .381 | .435 |
| Baseline Cortisol | Pain Tolerance | .123 | .160 | .256 |
| Baseline Alpha-amylase | Pain Threshold | .076 | .358 | .435 |
| Baseline Alpha-amylase | Pain Tolerance | .117 | .159 | .256 |
| Baseline Pulse | Pain Threshold | .154 | .062 | .198 |
| Baseline Pulse | Pain Tolerance | -.006 | .074 | .938 |
| Baseline MAP | Pain Threshold | .179 | **.030** | .198 |
| Baseline MAP | Pain Tolerance | .147 | .074 | .198 |

**Supplementary Table 5. Full table of moderation analyses.** *ß = estimate, CI = Confidence Interval, $f^2$ = Cohen's $f^2$ effect size, FPQ = Fear of Pain Questionnaire, LL = Lower Limit, MAP = Mean Arterial Pressure, n = Sample Size, NA = not applicable, p(-FDR) = Probability value (adjusted for multiple comparisons using False Discovery Rate correction), PCS = Pain Catastrophizing Scale, $R^2$ = coefficient of determination, SE = Standard Error, t = Test Statistic, TSK = Tampa Scale of Kinesiophobia, UL = Upper Limit. *Bootstrapped when assumptions were not met.*

| Outcome | Predictor | Moderator | Effect | *B* (unstd.) | SE | *t* | 95% CI LL | 95% CI UL | *β* (std.) | *p* | *p* (-FDR) | $\Delta R^2$ | $R^2$ | $f^2$ | *n* |
|---|---|---|---|---|---|---|---|---|---|---|---|---|---|---|---|
| Δ Pain threshold | Δ MAP | PCS sum score | Predictor | -0.057 | 0.034 | -1.671 | -0.125 | 0.011 | -0.203 | .099 | NA | NA | 0.056 | NA | 73 |
| Δ Pain threshold | Δ MAP | PCS sum score | Moderator | 0.029 | 0.034 | 0.840 | -0.040 | 0.097 | 0.100 | .404 | NA | NA | 0.056 | NA | 73 |
| Δ Pain threshold | Δ MAP | PCS sum score | Interaction | -0.003 | 0.005 | -0.605 | -0.012 | 0.006 | -0.072 | .547 | .915 | 0.005 | 0.056 | 0.005 | 73 |
| Δ Pain threshold | Δ MAP | TSK sum score | Predictor | -0.056 | 0.032 | -1.739 | -0.121 | 0.008 | -0.202 | .086 | NA | NA | 0.090 | NA | 73 |
| Δ Pain threshold | Δ MAP | TSK sum score | Moderator | 0.081 | 0.049 | 1.667 | -0.016 | 0.178 | 0.196 | .100 | NA | NA | 0.090 | NA | 73 |
| Δ Pain threshold | Δ MAP | TSK sum score | Interaction | -0.008 | 0.007 | -1.193 | -0.021 | 0.005 | -0.142 | .237 | .711 | 0.019 | 0.090 | 0.021 | 73 |
| Δ Pain threshold | Δ MAP | FPQ sum score | Predictor | -0.043 | 0.035 | -1.217 | -0.114 | 0.028 | -0.154 | .228 | NA | NA | 0.071 | NA | 73 |
| Δ Pain threshold | Δ MAP | FPQ sum score | Moderator | 0.018 | 0.018 | 0.972 | -0.019 | 0.054 | 0.115 | .335 | NA | NA | 0.071 | NA | 73 |
| Δ Pain threshold | Δ MAP | FPQ sum score | Interaction | -0.003 | 0.003 | -1.200 | -0.009 | 0.002 | -0.157 | .234 | .711 | 0.019 | 0.071 | 0.021 | 73 |
| Δ Pain threshold | Δ Pulse | PCS sum score | Predictor | -0.003 | 0.015 | -0.196 | -0.033 | 0.027 | -0.025 | .845 | NA | NA | 0.004 | NA | 73 |
| Δ Pain threshold | Δ Pulse | PCS sum score | Moderator | 0.017 | 0.035 | 0.480 | -0.053 | 0.087 | 0.059 | .632 | NA | NA | 0.004 | NA | 73 |
| Δ Pain threshold | Δ Pulse | PCS sum score | Interaction | 0.000 | 0.003 | 0.125 | -0.005 | 0.005 | 0.019 | .901 | .915 | 0.000 | 0.004 | 0.000 | 73 |
| Δ Pain threshold | Δ Pulse | TSK sum score | Predictor | -0.006 | 0.015 | -0.404 | -0.037 | 0.025 | -0.053 | .688 | NA | NA | 0.025 | NA | 73 |
| Δ Pain threshold | Δ Pulse | TSK sum score | Moderator | 0.066 | 0.050 | 1.323 | -0.034 | 0.166 | 0.161 | .190 | NA | NA | 0.025 | NA | 73 |
| Δ Pain threshold | Δ Pulse | TSK sum score | Interaction | -0.000 | 0.003 | -0.107 | -0.007 | 0.006 | -0.015 | .915 | .915 | 0.000 | 0.025 | 0.000 | 73 |
| Δ Pain threshold | Δ Pulse | FPQ sum score | Predictor | -0.006 | 0.015 | -0.387 | -0.036 | 0.024 | -0.050 | .700 | NA | NA | 0.008 | NA | 73 |
| Δ Pain threshold | Δ Pulse | FPQ sum score | Moderator | 0.013 | 0.019 | 0.670 | -0.025 | 0.051 | 0.082 | .505 | NA | NA | 0.008 | NA | 73 |
| Δ Pain threshold | Δ Pulse | FPQ sum score | Interaction | -0.000 | 0.001 | -0.271 | -0.002 | 0.002 | -0.032 | .787 | .915 | 0.001 | 0.008 | 0.001 | 73 |
| Δ Pain threshold | Δ Alpha-amylase | PCS sum score | Predictor | -0.006 | 0.004 | -1.345 | -0.015 | 0.003 | -0.157 | .183 | NA | NA | 0.104 | NA | 72 |
| Δ Pain threshold | Δ Alpha-amylase | PCS sum score | Moderator | -0.005 | 0.034 | -0.146 | -0.073 | 0.063 | -0.017 | .884 | NA | NA | 0.104 | NA | 72 |
| Δ Pain threshold | Δ Alpha-amylase | PCS sum score | Interaction | -0.002* | 0.001 | -2.605 | -0.004* | -0.000* | -0.400 | **.017*** | .072 | 0.089 | 0.104 | 0.100 | 72 |
| Δ Pain threshold | Δ Alpha-amylase | TSK sum score | Predictor | -0.002 | 0.005 | -0.438 | -0.012 | 0.008 | -0.056 | .663 | NA | NA | 0.025 | NA | 72 |
| Δ Pain threshold | Δ Alpha-amylase | TSK sum score | Moderator | 0.046 | 0.053 | 0.870 | -0.060 | 0.152 | 0.112 | .388 | NA | NA | 0.025 | NA | 72 |
| Δ Pain threshold | Δ Alpha-amylase | TSK sum score | Interaction | 0.000 | 0.001 | 0.337 | -0.001 | 0.002 | 0.035 | .737 | .915 | 0.002 | 0.025 | 0.002 | 72 |
| Δ Pain threshold | Δ Alpha-amylase | FPQ sum score | Predictor | -0.003 | 0.004 | -0.741 | -0.012 | 0.005 | -0.086 | .461 | NA | NA | 0.108 | NA | 72 |
| Δ Pain threshold | Δ Alpha-amylase | FPQ sum score | Moderator | 0.009 | 0.018 | 0.501 | -0.027 | 0.045 | 0.058 | .618 | NA | NA | 0.108 | NA | 72 |

| | | | | | | | | | | | | | | |
|---|---|---|---|---|---|---|---|---|---|---|---|---|---|---|
| Δ Pain threshold | Δ Alpha-amylase | FPQ sum score | Interaction | -0.001 | 0.000 | -2.583 | -0.002 | -0.000 | -0.292 | **.012** | .072 | 0.087 | 0.108 | 0.098 | 72 |
| Δ Pain threshold | Δ Cortisol | PCS sum score | Predictor | -0.004 | 0.040 | -0.097 | -0.085 | 0.077 | -0.012 | .923 | NA | NA | 0.010 | NA | 68 |
| Δ Pain threshold | Δ Cortisol | PCS sum score | Moderator | -0.011 | 0.035 | -0.302 | -0.081 | 0.060 | -0.038 | .763 | NA | NA | 0.010 | NA | 68 |
| Δ Pain threshold | Δ Cortisol | PCS sum score | Interaction | 0.005 | 0.007 | 0.751 | -0.009 | 0.019 | 0.114 | .455 | .911 | 0.009 | 0.010 | 0.009 | 68 |
| Δ Pain threshold | Δ Cortisol | TSK sum score | Predictor | -0.002 | 0.041 | -0.048 | -0.084 | 0.080 | -0.006 | .962 | NA | NA | 0.016 | NA | 68 |
| Δ Pain threshold | Δ Cortisol | TSK sum score | Moderator | 0.047 | 0.049 | 0.955 | -0.051 | 0.144 | 0.119 | .343 | NA | NA | 0.016 | NA | 68 |
| Δ Pain threshold | Δ Cortisol | TSK sum score | Interaction | 0.004 | 0.010 | 0.416 | -0.015 | 0.023 | 0.064 | .679 | .915 | 0.003 | 0.016 | 0.003 | 68 |
| Δ Pain threshold | Δ Cortisol | FPQ sum score | Predictor | -0.003 | 0.040 | -0.075 | -0.083 | 0.077 | -0.009 | .940 | NA | NA | 0.016 | NA | 68 |
| Δ Pain threshold | Δ Cortisol | FPQ sum score | Moderator | 0.007 | 0.018 | 0.379 | -0.030 | 0.044 | 0.047 | .706 | NA | NA | 0.016 | NA | 68 |
| Δ Pain threshold | Δ Cortisol | FPQ sum score | Interaction | -0.003 | 0.003 | -0.973 | -0.010 | 0.004 | -0.138 | .334 | .802 | 0.015 | 0.016 | 0.015 | 68 |
| cortisol stress responders only | | | | | | | | | | | | | | | |
| Δ Pain threshold | Δ Cortisol | PCS sum score | Predictor | 0.036 | 0.056 | 0.648 | -0.076 | 0.149 | 0.095 | .520 | NA | NA | 0.024 | NA | 50 |
| Δ Pain threshold | Δ Cortisol | PCS sum score | Moderator | 0.003 | 0.039 | 0.081 | -0.076 | 0.082 | 0.012 | .936 | NA | NA | 0.024 | NA | 50 |
| Δ Pain threshold | Δ Cortisol | PCS sum score | Interaction | 0.007 | 0.008 | 0.797 | -0.010 | 0.024 | 0.135 | .429 | NA | 0.013 | 0.024 | 0.014 | 50 |
| Δ Pain threshold | Δ Cortisol | TSK sum score | Predictor | 0.034 | 0.057 | 0.598 | -0.080 | 0.148 | 0.089 | .553 | NA | NA | 0.027 | NA | 50 |
| Δ Pain threshold | Δ Cortisol | TSK sum score | Moderator | 0.049 | 0.057 | 0.860 | -0.066 | 0.164 | 0.129 | .394 | NA | NA | 0.027 | NA | 50 |
| Δ Pain threshold | Δ Cortisol | TSK sum score | Interaction | 0.006 | 0.013 | 0.440 | -0.021 | 0.032 | 0.080 | .662 | NA | 0.004 | 0.027 | 0.004 | 50 |
| Δ Pain threshold | Δ Cortisol | FPQ sum score | Predictor | 0.041 | 0.056 | 0.724 | -0.072 | 0.154 | 0.106 | .473 | NA | NA | 0.018 | NA | 50 |
| Δ Pain threshold | Δ Cortisol | FPQ sum score | Moderator | -0.008 | 0.021 | -0.392 | -0.050 | 0.034 | -0.058 | .697 | NA | NA | 0.018 | NA | 50 |
| Δ Pain threshold | Δ Cortisol | FPQ sum score | Interaction | 0.001* | 0.004 | 0.388 | -0.014* | 0.129* | 0.062 | .852* | NA | 0.003 | 0.018 | 0.003 | 50 |

| | | | | | | | | | | | | | | | |
|---|---|---|---|---|---|---|---|---|---|---|---|---|---|---|---|
| Δ Pain tolerance | Δ MAP | PCS sum score | Predictor | -0.016 | 0.012 | -1.322 | -0.040 | 0.008 | -0.163 | .191 | NA | NA | 0.033 | NA | 73 |
| Δ Pain tolerance | Δ MAP | PCS sum score | Moderator | 0.011 | 0.012 | 0.877 | -0.013 | 0.035 | 0.106 | .383 | NA | NA | 0.033 | NA | 73 |
| Δ Pain tolerance | Δ MAP | PCS sum score | Interaction | 0.001* | 0.002 | 0.288 | -0.003* | 0.003* | 0.035 | .676* | .900 | 0.001 | 0.033 | 0.001 | 73 |
| Δ Pain tolerance | Δ MAP | TSK sum score | Predictor | -0.011 | 0.012 | -0.996 | -0.035 | 0.012 | -0.119 | .323 | NA | NA | 0.043 | NA | 73 |
| Δ Pain tolerance | Δ MAP | TSK sum score | Moderator | 0.005 | 0.017 | 0.317 | -0.029 | 0.040 | 0.038 | .752 | NA | NA | 0.043 | NA | 73 |
| Δ Pain tolerance | Δ MAP | TSK sum score | Interaction | -0.003* | 0.002 | -1.304 | -0.009* | 0.000* | -0.160 | .103* | .900 | 0.024 | 0.043 | 0.025 | 73 |
| Δ Pain tolerance | Δ MAP | FPQ sum score | Predictor | -0.015 | 0.012 | -1.241 | -0.040 | 0.009 | -0.158 | .219 | NA | NA | 0.055 | NA | 73 |
| Δ Pain tolerance | Δ MAP | FPQ sum score | Moderator | 0.010 | 0.006 | 1.564 | -0.003 | 0.023 | 0.186 | .122 | NA | NA | 0.055 | NA | 73 |
| Δ Pain tolerance | Δ MAP | FPQ sum score | Interaction | 0.000* | 0.001 | 0.157 | -0.001* | 0.002* | 0.021 | .853* | .900 | 0.000 | 0.055 | 0.000 | 73 |
| Δ Pain tolerance | Δ Pulse | PCS sum score | Predictor | 0.001 | 0.005 | 0.244 | -0.009 | 0.012 | 0.031 | .808 | NA | NA | 0.012 | NA | 73 |
| Δ Pain tolerance | Δ Pulse | PCS sum score | Moderator | 0.009 | 0.012 | 0.749 | -0.015 | 0.033 | 0.091 | .457 | NA | NA | 0.012 | NA | 73 |
| Δ Pain tolerance | Δ Pulse | PCS sum score | Interaction | -0.000* | 0.001 | -0.379 | -0.002* | 0.001* | -0.058 | .475* | .900 | 0.002 | 0.012 | 0.002 | 73 |
| Δ Pain tolerance | Δ Pulse | TSK sum score | Predictor | 0.002 | 0.005 | 0.270 | -0.009 | 0.012 | 0.035 | .788 | NA | NA | 0.005 | NA | 73 |
| Δ Pain tolerance | Δ Pulse | TSK sum score | Moderator | -0.001 | 0.018 | -0.055 | -0.036 | 0.034 | -0.007 | .957 | NA | NA | 0.005 | NA | 73 |
| Δ Pain tolerance | Δ Pulse | TSK sum score | Interaction | -0.001* | 0.001 | -0.410 | -0.002* | 0.002* | -0.060 | .611* | .900 | 0.002 | 0.005 | 0.002 | 73 |
| Δ Pain tolerance | Δ Pulse | FPQ sum score | Predictor | 0.001 | 0.005 | 0.169 | -0.009 | 0.011 | 0.021 | .867 | NA | NA | 0.033 | NA | 73 |
| Δ Pain tolerance | Δ Pulse | FPQ sum score | Moderator | 0.009 | 0.006 | 1.462 | -0.004 | 0.023 | 0.177 | .148 | NA | NA | 0.033 | NA | 73 |
| Δ Pain tolerance | Δ Pulse | FPQ sum score | Interaction | 0.000* | 0.000 | 0.127 | -0.000* | 0.001* | 0.015 | .857* | .900 | 0.000 | 0.033 | 0.000 | 73 |

| | | | | | | | | | | | | | | | |
|---|---|---|---|---|---|---|---|---|---|---|---|---|---|---|---|
| Δ Pain tolerance | Δ Alpha-amylase | PCS sum score | Predictor | 0.002 | 0.002 | 1.073 | -0.002 | 0.005 | 0.131 | .287 | NA | NA | 0.025 | NA | 72 |
| Δ Pain tolerance | Δ Alpha-amylase | PCS sum score | Moderator | 0.009 | 0.012 | 0.729 | -0.016 | 0.034 | 0.091 | .469 | NA | NA | 0.025 | NA | 72 |
| Δ Pain tolerance | Δ Alpha-amylase | PCS sum score | Interaction | 0.000* | 0.000 | 0.359 | -0.000* | 0.001* | 0.057 | .578* | .900 | 0.002 | 0.025 | 0.002 | 72 |
| Δ Pain tolerance | Δ Alpha-amylase | TSK sum score | Predictor | 0.002 | 0.002 | 1.058 | -0.002 | 0.005 | 0.136 | .294 | NA | NA | 0.020 | NA | 72 |
| Δ Pain tolerance | Δ Alpha-amylase | TSK sum score | Moderator | 0.007 | 0.019 | 0.386 | -0.030 | 0.044 | 0.050 | .701 | NA | NA | 0.020 | NA | 72 |
| Δ Pain tolerance | Δ Alpha-amylase | TSK sum score | Interaction | -0.000* | 0.000 | -0.309 | -0.000* | 0.000* | -0.033 | .598* | .900 | 0.001 | 0.020 | 0.001 | 72 |
| Δ Pain tolerance | Δ Alpha-amylase | FPQ sum score | Predictor | 0.002 | 0.002 | 1.025 | -0.002 | 0.005 | 0.122 | .309 | NA | NA | 0.055 | NA | 72 |
| Δ Pain tolerance | Δ Alpha-amylase | FPQ sum score | Moderator | 0.008 | 0.006 | 1.263 | -0.005 | 0.021 | 0.151 | .211 | NA | NA | 0.055 | NA | 72 |
| Δ Pain tolerance | Δ Alpha-amylase | FPQ sum score | Interaction | -0.000* | 0.000 | -0.876 | -0.000* | 0.000* | -0.102 | .120* | .900 | 0.011 | 0.055 | 0.011 | 72 |
| Δ Pain tolerance | Δ Cortisol | PCS sum score | Predictor | 0.018 | 0.013 | 1.378 | -0.008 | 0.043 | 0.169 | .173 | NA | NA | 0.051 | NA | 68 |
| Δ Pain tolerance | Δ Cortisol | PCS sum score | Moderator | 0.013 | 0.011 | 1.187 | -0.009 | 0.035 | 0.145 | .240 | NA | NA | 0.051 | NA | 68 |
| Δ Pain tolerance | Δ Cortisol | PCS sum score | Interaction | 0.001* | 0.002 | 0.604 | -0.004* | 0.007* | 0.089 | .648* | .900 | 0.005 | 0.051 | 0.006 | 68 |
| Δ Pain tolerance | Δ Cortisol | TSK sum score | Predictor | 0.012 | 0.013 | 0.921 | -0.014 | 0.037 | 0.113 | .361 | NA | NA | 0.067 | NA | 68 |
| Δ Pain tolerance | Δ Cortisol | TSK sum score | Moderator | -0.007 | 0.015 | -0.478 | -0.038 | 0.023 | -0.058 | .634 | NA | NA | 0.067 | NA | 68 |
| Δ Pain tolerance | Δ Cortisol | TSK sum score | Interaction | 0.005* | 0.003 | 1.611 | -0.001* | 0.012* | 0.240 | .144* | .900 | 0.038 | 0.067 | 0.041 | 68 |
| Δ Pain tolerance | Δ Cortisol | FPQ sum score | Predictor | 0.018 | 0.012 | 1.444 | -0.007 | 0.042 | 0.171 | .154 | NA | NA | 0.110 | NA | 68 |
| Δ Pain tolerance | Δ Cortisol | FPQ sum score | Moderator | 0.013 | 0.006 | 2.349 | 0.002 | 0.025 | 0.279 | **.022** | NA | NA | 0.110 | NA | 68 |
| Δ Pain tolerance | Δ Cortisol | FPQ sum score | Interaction | -0.001* | 0.001 | -0.980 | -0.004* | 0.001* | -0.133 | .409* | .900 | 0.013 | 0.110 | 0.015 | 68 |
| cortisol stress responders only | | | | | | | | | | | | | | | |
| Δ Pain tolerance | Δ Cortisol | PCS sum score | Predictor | -0.009 | 0.012 | -0.767 | -0.034 | 0.015 | -0.108 | .447 | NA | NA | 0.101 | NA | 50 |
| Δ Pain tolerance | Δ Cortisol | PCS sum score | Moderator | 0.016 | 0.009 | 1.795 | -0.002 | 0.033 | 0.258 | .079 | NA | NA | 0.101 | NA | 50 |
| Δ Pain tolerance | Δ Cortisol | PCS sum score | Interaction | -0.001* | 0.002 | -0.558 | -0.007* | 0.002* | -0.091 | .628* | NA | 0.006 | 0.101 | 0.007 | 50 |
| Δ Pain tolerance | Δ Cortisol | TSK sum score | Predictor | -0.014 | 0.013 | -1.043 | -0.040 | 0.013 | -0.155 | .302 | NA | NA | 0.026 | NA | 50 |
| Δ Pain tolerance | Δ Cortisol | TSK sum score | Moderator | 0.004 | 0.013 | 0.282 | -0.023 | 0.030 | 0.042 | .779 | NA | NA | 0.026 | NA | 50 |
| Δ Pain tolerance | Δ Cortisol | TSK sum score | Interaction | 0.002* | 0.003 | 0.509 | -0.002* | 0.006* | 0.093 | .378* | NA | 0.005 | 0.026 | 0.006 | 50 |
| Δ Pain tolerance | Δ Cortisol | FPQ sum score | Predictor | -0.011 | 0.013 | -0.857 | -0.036 | 0.015 | -0.123 | .396 | NA | NA | 0.060 | NA | 50 |
| Δ Pain tolerance | Δ Cortisol | FPQ sum score | Moderator | 0.006 | 0.005 | 1.341 | -0.003 | 0.016 | 0.193 | .187 | NA | NA | 0.060 | NA | 50 |
| Δ Pain tolerance | Δ Cortisol | FPQ sum score | Interaction | 0.001* | 0.001 | 0.576 | -0.001* | 0.002* | 0.090 | .439* | NA | 0.007 | 0.060 | 0.007 | 50 |
